# Unsupervised Idealization of Nano-Electronic Sensors Recordings with Concept Drifts: A Compressive Feature Learning Approach for Non-Stationary Single-Molecule Data Analysis

**DOI:** 10.1101/2020.05.02.074013

**Authors:** Mohamed Ouqamra

## Abstract

Single-molecule nanocircuits based on field-effect transistors (smFETs) are emerging and promising nano-bioelectronic sensors for the functional detection of molecular dynamics involved in biochemical transformations, in particular for applications in cancer thanks to a potentially better understanding of some hidden and complex molecular interactions. In fact, functionalized carbon nanotubes have been recently exploited to probe molecular events occurring at a single molecule scale with ultra high sensitivity and specificity, such as nucleic acids hybridization, enzyme folding in catalysis reactions, or protein-nucleic acids interactions. Extracting the kinetics and thermodynamics from such single-molecule dynamics implies robust analytic tools that can handle the complexity of the sensed reaction system changing between transient and steady-state molecular conformations, but also some challenging signal specificities, such as the multi-source composition of the recorded signals, the mixed and high-level noises, and the sensor baseline drift, leading to non-stationary time series. We present a new smFET data analysis framework, based on a compressive feature learning scheme to optimize unsupervised idealization of smFET traces, by a precise and accurate molecular events detection and states characterization algorithm, tailored for non-stationary signals at high sampling rate and long acquisition periods, without any prior knowledge on the data generating process nor signal pre-filtering. Experimental results show the accuracy and robustness of our trace idealization algorithm to stochastic state-space models, and better performances than commonly used hidden Markov models.

## Introduction

Single-molecule experiments present a great interest to analyze the structural and interactions dynamics of complex molecules such as biological macro-molecules. Fluorescence-based techniques (total internal reflection fluorescence microscope, confocal microscopy and fluorescence correlation spectroscopy), opposed or combined to force-based techniques (optical trap, magnetic trap and atomic force microscope) (1–4), offer the possibility to study molecules one at a time and to highlight reaction pathways and reaction intermediates that ensemble methods tend to hide because of their effect of averaging (5).

More recently, another approach has been emerging based on single-molecule field-effect transistors (smFETs), in which an individual molecule is immobilized on a nanoscale electrical circuit, such as a carbon nanotube (6–12) or silicon nanowire (13–16). By recording fluctuations in the electrical conductance of such circuits, studies have reported the real-time monitoring of transitions between different conformational states, such as hybridization (17–20), folding events in nucleic acids (17, 21), enzymatic catalysis with a ultra-high sensitivity and specificity (22, 23), and also other applications in biology and medicine, such as imaging (24, 25) and drug delivery (26, 27) to cancer and brain.

Detecting and modeling the kinetics and thermodynamics of such molecular interactions from smFET recordings require robust data analysis tools that can handle challenging signal specificities: 1) the stochastic nature of the biomolecular system, 2) the possible non-stationarity of the molecular dynamics of the reaction system, such as changes between transient and steady-state conformations, 3) the multi-source composition of the sensor response, aggregating all the contributions from the biochemical system with those of the measurement medium and the sensor components into a single output, 4) the mixed noises (AWGN, flicker, and impulse) characteristic of FET devices, 5) the sensor baseline drift that can occur during long acquisitions, and 6) the sizable amount of data generated by such recordings, all together resulting in complex time series to idealize into a state trajectory.

Several single-molecule data (SMD) analysis techniques have been developed for trace idealization, such as visual inspection (28), manual thresholding (29), or Markovian model-based approaches. In particular, hidden Markov models (HMMs) (30, 31) have been widely used for the analysis of fluorescence resonance energy transfer (FRET) time trajectories (32, 33), biological sequences in genomics (34),or protein modeling (35), but require restrictive assumptions and a priori knowledge of the likely kinetics model, which are hardly available in real conditions. In particular, the molecular state-space size needs to be predefined through initial guesses to prime the Baum-Welch learning algorithm (36), while the assumed markovianity of the probed conformational states remains to be verified a posteriori. More recently, a Bayesian non-parametric (BNP) implementation as infinite HMM, has been proposed (37) to overcome such drawback through a built-in model complexity adaptation to new incoming data. Nevertheless, this alternative approach has actually a context dependency because of the user-preset (hyper) priors related to the data generating process that must be carefully formulated for both the molecular reaction system itself, and the physics involved in the measurement acquisition chain, to avoid overfitting and misinterpretation. Furthermore, if BNP outperforms finite HMMs via the adaptive state space size to new data, their main limitation is to find the optimal beam sampling rate that converges to stationary chains, in other words the number of samples (MCMC iterations) to draw from the proposal distribution before reaching the true posterior is not learned from the data, despite some convergence diagnostics (38, 39) that can help in tuning the stop sampling criterion. Another approach based on the minimum description length (MDL) principle has been proposed for the idealization of ion channel recordings (40). Due to the robustness of MDL to non-stationary signals and its fast computational time, we derived this model-free approach to the smFET traces idealization through an entropy-based objective.

In this contribution, we propose a novel approach based on machine learning for signal processing to retrieve the hidden dynamic network of the molecular transition states, without any user-defined priors on the involved kinetics or the sensor characteristics. Our method is a hybrid algorithm combining model-free signal compression and clustering (cc) techniques as an unsupervised data preparation, followed by an expectation-maximization (EM) refinement, both leading to a more robust smFET trace idealization algorithm, called cc-EM (compressed clustered EM), for non-stationary and heterogeneous signals. The main idea is that if data-driven modeling using machine learning techniques can supply knowledge-based models beyond their assumption limitations, the more reliable the data, the more accurate will be the resulting model. We show how each data preparation step of cc-EM contributes to highlight the inner structure and dimensionality of the data, even under very noisy environments, and facilitates the inference task of the EM step to better estimate the states trajectory, but also the molecular state-space size by a model selection algorithm as a decision aid-tool. Validation on a wide range of smFET signal parameters and kinetic scenarios shows better detection performances (RMSE boxplots, precision and recall scores through ROC curves, right prediction rate of the state space size) than the Baum-Welch and Viterbi algorithms in HMM methods, making this entropy-based and unsupervised smFET trace idealization method more accurate and robust to kinetics modeling of non-stationary and heterogeneous multi-state dynamics from evolving and drifting data.

The remainder of this paper is structured as follows. In section I we detail the problem statement of smFET trace idealization. The section II details the derivation of the objective functions used in the proposed framework, while section III describes the test results and the performance metrics to assess the reliability and robustness of the learning algorithm given several signal and kinetic parameters. Finally, the detailed results conclude to the out-performance of the proposed cc-EM algorithm to learn from non-stationary and drifting smFET traces, quantitative and qualitative kinetic features of a multi-state model, both under real conditions and challenging simulated molecular dynamics.

## Unsupervised Idealization of Non-Stationary smFET signals

### A. Problem statement

Based on the Transition State theory, we assume that the single-molecule dynamics can be modeled by a piecewise regression of the smFET signal, leading to an idealized trace made of contiguous segments of locally constant electrical intensity, separated by instantaneous transitions of the conductance. Thus, the underlying probed single-molecule interactions can be explained by a non-linear and time-variant oscillating system whose time trajectory can be approximated by discretization of the emissions of the visited hidden states.

Let *y* be an smFET signal of length *N* that can be decomposed into a finite and discrete sum of homogeneous, locally stationary and uncorrelated sub-signals, associated to contiguous segments of variable lengths and intensities over time. In other words, the signal *y* is piecewise stationary, but some features of the data generating process may change abruptly at unknown changepoint instants,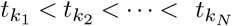. The problem of the retrospective detection of these breakpoints can be formulated as an iterative segmentation process of the signal *y*(*t*) into its idealized estimate *ŷ*(*t*), providing structural information through the temporal locations and the number *k* of abrupt changes, corresponding to the events or transitions that delimit over time some metastable molecular conformational states of the reaction system under monitoring, such as: 

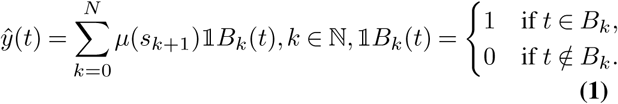

With *µ*(*s*_*k*+1_) the mean electrical conductance level of the *k* + 1^*th*^ segment induced by the *k*^*th*^ detected molecular event, while if no transition is detected (*k* = 0), the idealized trace *ŷ*(*t*) is comparable to the average intensity 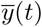 of the whole raw signal *y*(*t*), corresponding to a single state model of a thermodynamically stable reaction system. However, this trace idealization task can be complicated by some signal constraints that are characteristic of the smFET experiments, namely:

1. The non-stationarity of the recorded traces.
2. The measurement and process noises’ level that blur the states trajectory.
3. The signal distortion due to the sensor baseline drift over time that can locally induce fake states and transitions.
4. The multi-source composition of the sensor response to the molecular dynamics between the target and the receptor biomolecules functionalized on the surface of the transducer, according to a MISO (multi-input single output) configuration, leading to heterogeneous smFET signals *y*(*t*).

Such heterogeneity of the signal sources motivates the reformulation of the recorded observations *y*(*t*) at each time step *t*, as the sum of the emissions *e*(*t*) from the hidden molecular states *s*_*t*_, with the sensor baseline drift *d*(*t*), the measurement noise, *ϵ*(*t*), so that the resulting data generating model is: 

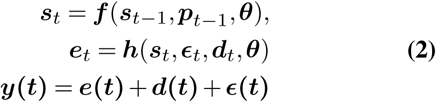

***f*** and ***h*** being respectively the unknown transition and emission matrices of a first-order Markov process, and ***p*** _*t−*1_ the process noise.

We aim to learn from the observations *y*(*t*) the underlying molecular dynamics through the states’ trajectory by the piecewise regression of the emissions *e*(*t*), leading to the idealized trace *ŷ*(*t*) defined i n e quation 1, t aking i nto account the above mentioned smFET signal constraints. The contributions of the proposed cc-EM framework are multiple:

1. cc-EM is a full automatic and unsupervised trace idealization method that does not require any precise definition of the data generating process, neither related to the underlying kinetics of the probed reaction system, nor to the physics of the measurements from the acquisition chain.
2. cc-EM can learn the states and rates under concept drift, while the resulting kinetic model can be updated as new incoming data are recorded.
3. Our framework includes an automatic baseline drift removal tool to avoid fake states and transitions in the resulting idealized trace.
4. cc-EM is robust to mixed noises (white Gaussian, flicker and impulse noise), even under weak SNR values (*SNR* = 1*dB*).
5. The cc-EM algorithm can handle complex asymmetrical transition probability matrices for Markov processes without detailed balance among the steady-states network, while it does not require any particular distribution for the states’ emissions to yield valid kinetic models.
6. No need of any analog or digital signal prefiltering (that could bias the kinetics estimates of the learned model) for cc-EM to achieve robust and reliable idealized traces.
7. The computational time *𝒪*(*n* log *n*) of the whole piece-wise segmentation process makes it compatible to large datasets mining.

### B. Proposed Approach

Our global methodology is defined in a 4-step approach for the analysis of smFET signals and the performance assessment of the proposed trace idealization algorithm:

1. Baseline Drift Removal
2. Trace Idealization
3. Kinetics Modeling
4. Transition Network Visualization

For the baseline drift removal stage, we implemented in the proposed framework a built-in drift compensation algorithm based on information theory (41), thus allowing to prepare the signal for an optimal trace idealization, since such parasitic drift may induce locally fake states and transitions leading in turn to erroneous kinetic models.

For the trace idealization stage, we propose a hybrid approach combining machine learning and statistical methods for signal processing. The cc-EM (compression-clustering-Expectation-Maximization) algorithm tackles the constraining imposition of a particular kinetic scheme and restrictive assumptions conditioning the validity of the outputs.

For the validation tests, unless otherwise specified in the corresponding experiment, for each kinetic and sensing parameter tested, we used 200 synthetic traces of 2500 data points. A synthetic trace generator allowing the user to experiment a wide range of smFET signals given different kinetic and sensing profiles is also implemented in the cc-EM framework. The smFET trajectories are generated in Matlab using a random walk process through a first-order Markov chain, given some tunable parameters enabling to simulate both the kinetics of the molecular dynamics through the states trajectories and the sensor’s response over time. The kinetic parameters include (1) the number of hidden states of the Markov process, (2) the initial probabilities of the hidden states if available or based on guesses, (3) the transition and emission matrices representing respectively the states’ life-time and the events rate, and the quantization step of the observations space, and (4) the concept drift corresponding to changes in the data generating process, for instance a sensor’s response that varies as the biochemical reaction under monitoring undergoes intermediates steps, or kinetic changes between the conformational states due to a catalyst. To simulate the observable effects of such reaction drifts, one can thus modulate the first and/or the second moment of the data distribution by concatenating several traces, each having different transition matrices, and/or different sizes representing the number of hidden states, that is a change in the molecular model itself.

The sensor’s response parameters include (1) the signal to noise ratio and the type of noise, additive white Gaussian, pink, shot and mixed noises, (2) a trend parasitic signal with an adjustable shape and oscillations amplitude, simulating the sensor’s baseline drift due to the slowly varying environmental conditions of the measurement medium, such as temperature, pressure, humidity, impacting the sensor’s response to the target analyte, (3) the sampling frequency and the acquisition duration that both define the length of the resulting smFET trace and the size of the dataset to process.

Finally, we compiled all the kinetics and thermodynamics parameters learned from the smFET recordings into a single-molecule data visualization tool to better highlight the underlying transition networks of the molecular conformational changes into an effective free energy landscape and reaction coordinates.

### C. cc-EM algorithm: an Entropy-based Piecewise Segmentation for Non-Stationary smFET Trace Idealization

We propose and detail the cc-EM algorithm, a compression and clustering signal preparation for Expectation-Maximization refinement, followed by a model selection, as a 4-step trace idealization method robust and tailored for non-stationary and heterogeneous single-molecule signals. The flowchart of the cc-EM algorithm is resumed in figure 1.

**Fig. 1.**
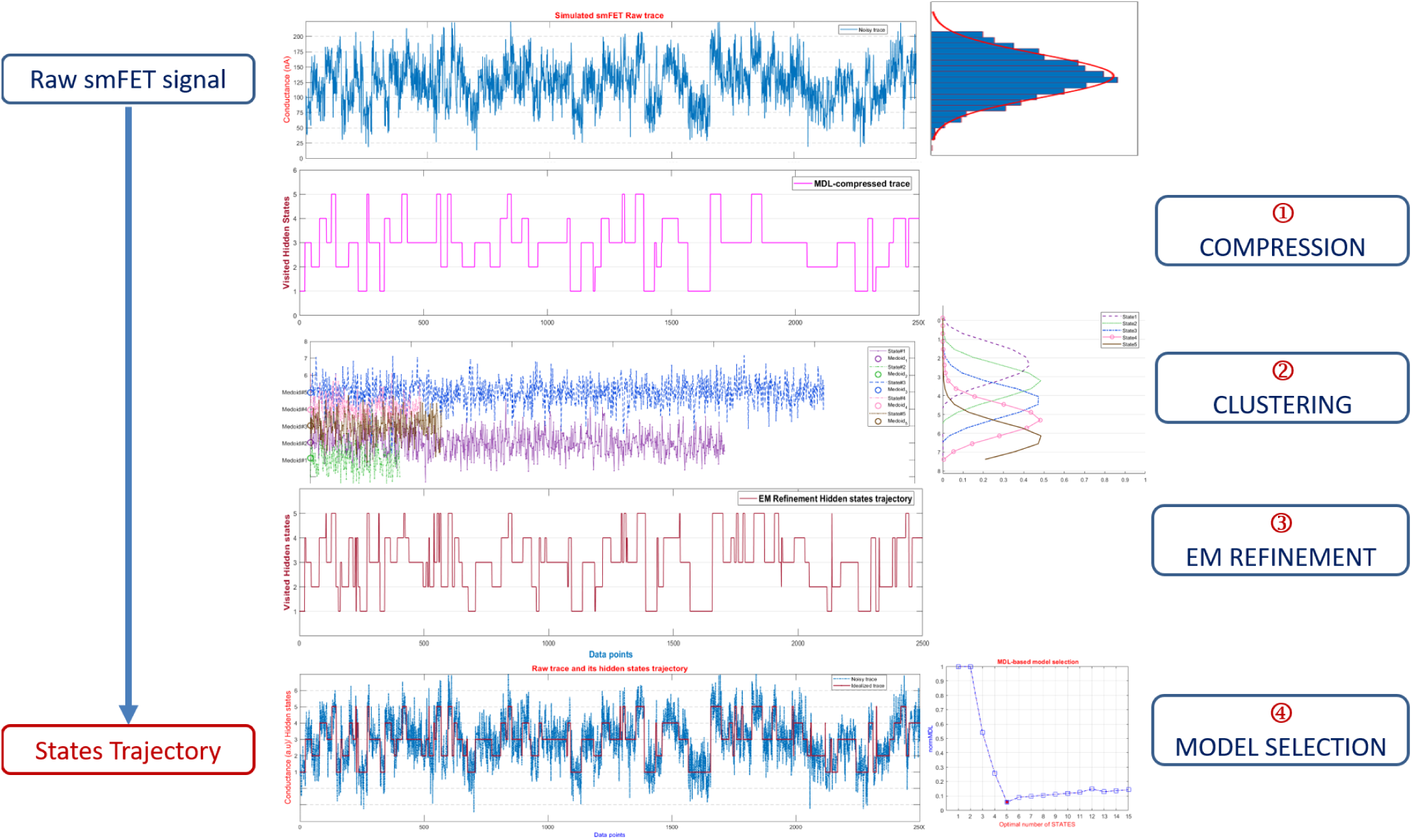
Flowchart of the cc-EM framework

<u>step 1: SIGNAL COMPRESSION</u>

Considering a smFET time series *Y* = {*y*_1_, *y*_2_, *…, y*_*N*_} of length *N*, to make the achieve a more robust trace idealization method to non-stationary and heterogeneous signals, we reformulated the MDL-based objective defined in (40) as a signal compression task according to: 

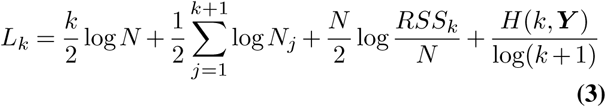

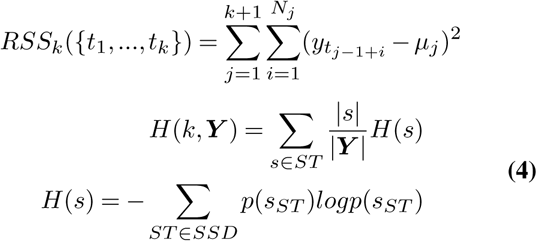

Such problem definition relies on information theory (42) and the principle that the more we compress the data, the more we have learned about them (43).The objective function *L*_*k*_ is the code-length in bits of the resulting state trajectory *ST*, so that the most likely idealized trace is the one that requires the less amount of bits to encode the state model parameters. The first term in Eq. 3 is the code-length in bits of the *k* abrupt concept drifts corresponding to the time locations of the change points between the states. The second term encodes the density population *N*_*j*_ of each segment or plateau *j* in the compressed trace, while we redefined the third term as the bit-length to encode the compression error between the piecewise trace and the raw signal. Minimizing the encoding cost of the molecular features according to these three first terms in Eq. 3, does not guaranty a lossless compression and a perfect retrieval of the hidden states trajectory. To achieve optimal event detection performances, we added an entropy term in the objective function *L*_*k*_, motivated by two main reasons.Firstly, because some molecular events can occur several times due to the biochemical reaction itself, but also to the multiple active sites functionalized on the transducer leading in either case to temporal or spatial redundancies in the recordings, while such events can have similar kinetics that can be assigned to a common state. Secondly, because of some outliers and aberrant values that can be recorded in the sensor’s response over time, and simulate fake sub-states. Thus, the last term 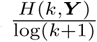, is the normalized entropy of the raw signal *Y* compressed and discretized into *k* + 1 discrete steps, and *H*(*k, Y*) being the entropy of the partitioned signal *Y* by each detected transition *k* into *k* + 1 plateaus of the resulting states trajectory *ST*, while |*s* denotes the density population of each state *s. H*(*s*) being the entropy of each detected state *s*, and *p*(*s*_*ST*_) the conditional probability of a data point in the signal *Y* to be allocated to the state *s* given the state trajectory *ST*, and given all other possible scenarios of states’ dwelling in the state space domain *SSD* According to the source coding literature (42, 44, 45) the entropy of a stochastic source measures the minimum average bit-length to compress the data generated by that source in a lossless manner. Ideally, one seeks to get a lossless signal compression that leads to a perfect match between the true hidden states trajectory and the compressed idealized trace, but depending on the noise level (1*dB ≤ SNR ≤* 5*dB*) and on the events rate of the molecular dynamics, the MDL-based idealization is most often a lossy compression resulting in missed states and transitions, and even to a wrong kinetic model, that is an erroneous state space size estimate. To overcome such drawback, our method consists in finding the optimal compression ratio corresponding to the optimal learning rate, according to the minimum description length principle for the parameters of a time-discrete finite state model. To this end, the entropy term in the compression cost function in equation 3 serves to penalize the possible redundancies in the raw signal that the two main limiting factors (noise level and event rate) tend to accentuate, so that the lower the entropy the closer we get to the optimal compression ratio, and to a lossless compressed states trajectory.

<u>step 2:CLUSTERING OF THE COMPRESSION PATTERNS</u>

Because the different probabilities *p*(*s*_*ST*_) involved in the compression objective function in 4 are not always available in real conditions, to reach the entropy lower bound (ELBO) that brings closer to a lossless compression, instead of using the maximum likelihood estimate of theses probabilities, we rather opted for a K-medoids (46) clustering of the different compression patterns obtained in step 1, to gather possible sub-states having some signal features similarities that are discovered from the data without supervision, into some common parent states associated here to each medoid of the partitioning. To do so, we implemented a swapping cost function that tests for each pair (*C*_*p*_, *S*_*j*_) of a potential true state *S*_*j*_, and a compression pattern *C*_*p*_ defined as the 4-tuple: (the length of the conductance state, the mean intensity of the state, the time location of the transition delimiting the current state from its neighbor, and the magnitude of such transition, which indirectly define the distances between the clusters), whether *C*_*p*_ is better than *S*_*j*_ as a final state (*C*_*p*_ and *S*_*j*_ being here associated with clusters), given the Mahalanobis distance *D*_*M*_ (47, 48) : 

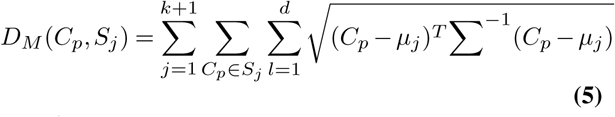

with *k*, the number of abrupt concept drift separating the *k* + 1 discrete piecewise segments, *d* the dimensionality of the data, (*d* = 4 for smFET time series, according to the definition of the compression pattern detailed above), and *µ*_*j*_, Σ, respectively the mean and the covariance matrix of the Gaussian distribution associated to the emissions of each state *s*_*j*_. The choice of such similarity metric relies on its higher robustness to outliers, its scale-invariance to multi-source signals, and on its consideration of the correlations in the processed signals. The effect of such second step is to lower the false positive rate of the molecular transitions detection task, since we get more stable clusters than the compressed plateaus obtained in step 1, leading to a better separation of the states even if their spectral emissions overlap in the raw signal. We take advantage from the first compression step to infer a cluster number range of [1, *sss*_*MDL*_], into which the compression patterns are allocated. *sss*_*MDL*_ being the number of unique plateaus in the MDL compressed trace, so that we assume *sss*_*MDL*_ to be the upper bound of the molecular state space size to scan, because already compressed according to 3, but not with the optimal compression ratio, resulting in a lossy compression of the states’ emissions and the overfitting of the number of states. Compared to the infinite HMM, our compression and clustering of the state space domain *sss* can be respectively seen as the Griffiths-EngenMcCloskey stick-breaking construction of the base probability for the Dirichlet process (49, 50), and the beam sampler to truncate the resulting potentially too large state space (51, 52).

We also compared the K-medoids partitioning efficiency to the K-means clustering (53, 54), using an Euclidean distance as similarity measure, stating that since the smFET conductance values follow an uncorrelated univariate normal distribution, we have a diagonal covariance matrix and the Mahalanobis distance reduces to the normalized Euclidean distance, which relatively limits possible bias in the performance comparison between the two partitioning methods. We compared the goodness of fit of the resulting idealized traces obtained by the objective function in (40), to the states trajectory estimate using the proposed 2-steps signal compression and clustering, either by the K-medoids or the K-means partitioning, given a varying additive white Gaussian noise (AWGN) level on 200 synthetic traces of 2500 data points each, simulating a 2-states first-order Markovian random walk. Figure 3 shows how the K-medoids clustering clearly improves the performances of the compression step alone, especially as the noise level increases (SNR<15dB), while it clearly outperforms the K-means clustering state trajectory estimates. We also compared the precision and recall scores between the two data preparation steps. The ROC curves in figure 3 highlights how the K-medoids clustering post MDL-based signal compression, with an *AUC* = 95.4% is a decisive step in improving the performances of the MDL-based event detection algorithm (*AUC* = 80%), and how it is also a better partitioning choice compared to the K-means outputs (*AUC* = 76%).

**Fig. 2.**
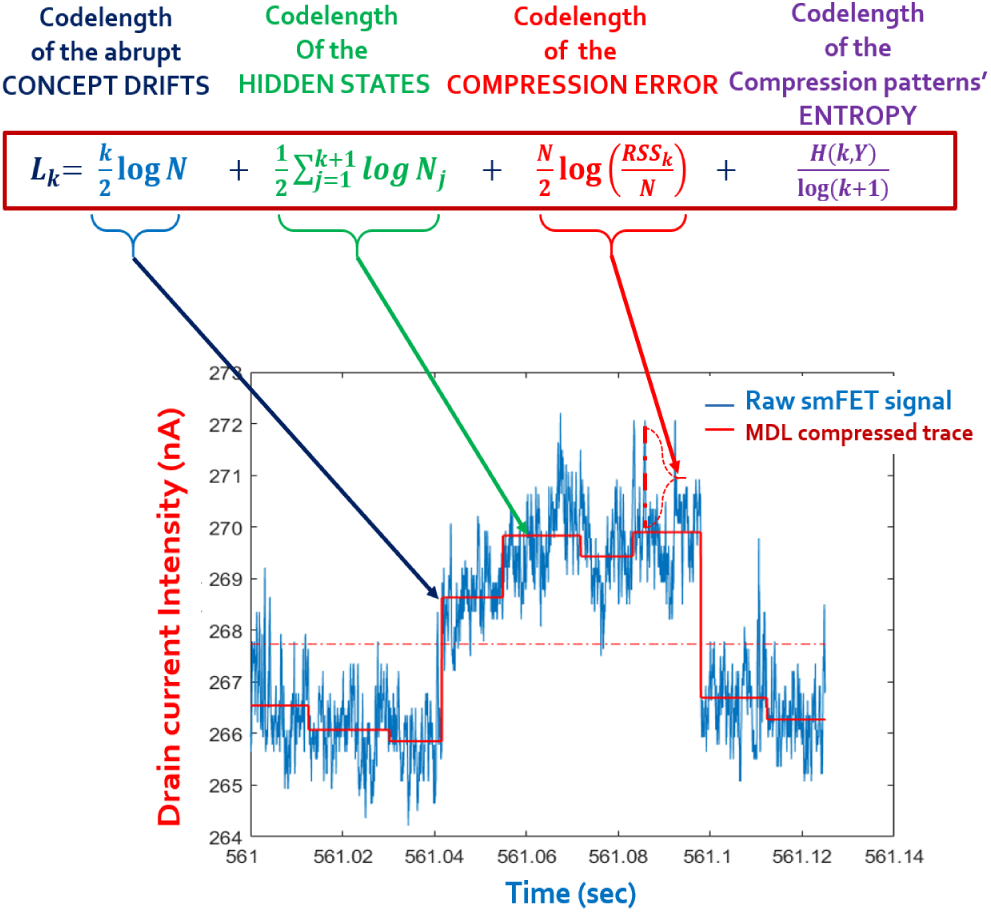
Illustration of the MDL-based signal compression objective function for the piecewise segmentation process of smFET trace idealization. The first two terms are the bit-length of the state-model parameters, the third term is to encode the compression error, and the last term is the code-length of the states’ entropy.

**Fig. 3.**
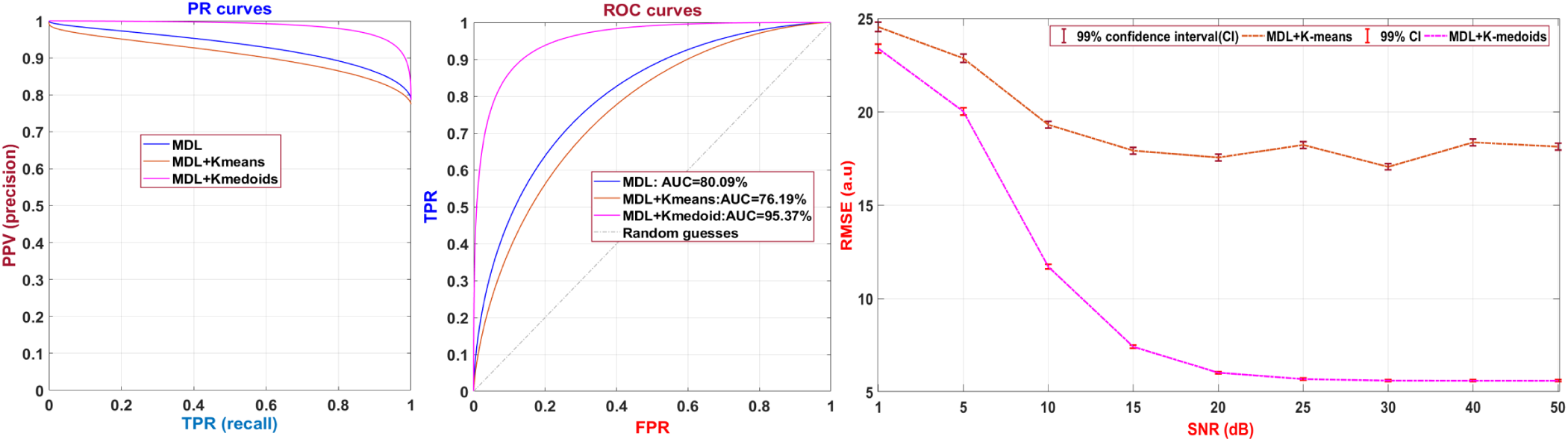
Performance comparison between the K-means and K-medoids clustering of the compression patterns in terms of: (left) Precision-Recall curves (middle) AUC of the ROC curves given a constant noise level of 1dB, and a number of hidden states varying between 2 and 15. Tests have been performed on 200 simulated samples of 2500 points each, using macro-averaging of the precision and recall scores calculated from the resulting confusion matrices for each test. (right) Goodness of fit through the residuals between the estimated idealized traces obtained by MDL+K-means or MDL+K-medoids and the true states trajectory of 200 synthetic traces of 2500 points each, simulating a 2-states first order Markov chain, corrupted by an additive white Gaussian noise (AWGN) with an increasing level (1*dB ≤ SNR ≤* 50*dB*).

<u>step 3: EXPECTATION-MAXIMIZATION REFINEMENT</u>

Once the signal has been compressed and clustered, depending on both the noise type (AWGN,pink, flicker, mixed) and level (*SNR ≥* 5*dB*), and the transitions frequency due to the molecular dynamics, these two steps may be sufficient to obtain accurate idealized traces. Nevertheless, in real conditions these critical sensing and kinetic parameters may require to infer some possible missed states and transitions through an expectation-maximization (EM) refinement scheme, as a third step.

The main added value of our approach is that by training the EM algorithm on the compressed and clustered piecewise trace, we initialize the learning algorithm with more accurate priors than random guesses, in the sense that the MLD+K-medoids preliminary state trajectory is much more closer to the ground truth than any other random starts, which also makes the cc-EM algorithm to faster converge to the optimal clustering solution, thanks to the known posterior distribution of each state given the data. More importantly, by feeding the EM algorithm with such prior states trajectory, we take advantage from the robustness of the MDL compression stage to non-stationary distributions, using it as a front-end data preprocessing so that we manage to bypass the restrictive homoscedasticity assumption of the EM algorithm. Finally, because the EM refinement requires the number of states to be a priori defined, we use the number of unique plateaus in the MDL+k-medoids output as initial guess of the lower bound of the state space size, and the number of unique plateaus in the MDL-based compressed trace as the upper bound, while the last step as a model selection will automatically choose the most likely states trajectory that best explains the data, wrt the best trade-off between model complexity and goodness of fit. Using the probabilities of each data point to be emitted by each state as priors for the EM refinement, enables better robustness to low SNR values, since it reduces the risk of data-to-cluster miss-assignments, and the under-fitting of the state space size, thanks to the higher sensitivity to local intensity fluctuations of the MDL-based compression stage that can separate states with very close signal intensity. We propose to derive the maximum a posteriori expectation-maximization (MAP-EM) steps (55) for learning the set of parameters 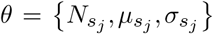 of a generalized smFET state-model, without any kinetic model assumption. In the particular case of Markovian processes the set of parameters *λ* ={Π, *A, B}*, (Π being the initial probabilities of each state, *A* the matrix of the transition probabilities between the states, and *B* the matrix of the emission probabilities) is learned by a Hidden Markov model using a modified forward-filtering backward sampling algorithm adapted for continuous-valued emissions, and applied on each states sequence sampled from its corresponding compressed and clustered trace of the screened state space, while the most likely path is deduced by the Viterbi algorithm (56)

**Table 1.** Notation Summary

| Variable Notation | Designation |
| --- | --- |
| $N$ | Sample size (number of observation) |
| $Y$ | Random measurement vector |
| $y \in \mathbb{R}^d$ | Given observation, $y$ is a realization of $Y$ |
| $i$ | Observation index |
| $j$ | State index |
| $k$ | Transition index |
| $s_j$ | $j^{th}$ detected state |
| $\mu_{s_j}$ | Mean intensity of the piecewise segment allocated to the $j^{th}$ state |
| $N_{s_j}$ | Relative population density of the $j^{th}$ state |
| $sss$ | State space size, i.e. the number of states |
| $\sigma_{s_j}$ | Noise variance of the $j^{th}$ state |
| $\theta \in \Omega$ | Parameters to estimate of the state-model; $\Omega$ being the parameter space |
| $\theta^m$ | $m^{th}$ estimate of $\theta$ |
| $p(y\vert \theta)$ | Density of $y$ given $\theta$: $p(Y = y_i\vert \theta)$ |
| $p_{y_i s_j}$ | probability of the $i^{th}$ observation to belong to the $j^{th}$ detected state |

As mentioned above, we can extract several information from the MDL-compressed and the K-medoids clustered traces:

1. let *sss* be the likelihood of the state space size, with *sss*_*min*_ and *sss*_*max*_ respectively the lower and upper bound of *sss*, deduced from the number of detected states at the clustering and the lossy compression stages respectively, while if this number is the same for these two data preparation steps, we expand the state space domain to explore of one state *sss* = *sss* ± 1 to avoid potential underfitting or overfitting.
2. Among the outputs of the k-medoids algorithm, we can access to all the distances *d*(*y*_*i*_, *m*_*j*_) between each observation *y*_*i*_ and each medoid *m*_*j*_ representing here the mean intensity of each detected state *s*_j_. To transform these distances into probabilities, we apply the softmax function on each row vector *z* = *d*(*y, m*), *i ∈*[1, *N*], *j* ∈ [1, *sss*_*max*_]:

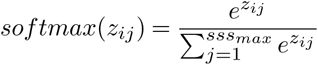

For each states trajectory of the state space domain, we apply the MAP-EM algorithm derivation to the smFET idealized trace refinement that runs as follows:

#### 1. Initialization

Let the number of iteration *m* = 0, and make an initial estimate *θ*^*m*^ for *θ*, using the following priors: 

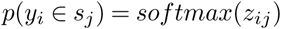

#### 2. Expectation

Given the observed data *y*_*i*_ and assuming for the moment that the current guess *θ*^*m*^ is correct, formulate the conditional probability distribution *p*(*x|y*_*i*_, *θ*^*m*^) for the complete data *x* = (observed variables,latent variables), which is the probability that the sequence of hidden states has really generated the recorded sample *Y* of *N* observations (if we consider the maximum likelihood estimate), or more precisely the latent state *s*_*j*_ from which the observation *y*_*i*_ originates.

Using the conditional probability distribution *p*(*x|y*_*i*_, *θ*^*m*^), define the conditional expected log-likelihood, also called the Q-function: 

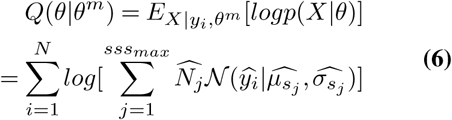

For each *j*^*th*^ detected segment, the maximum likelihood estimates (MLEs) for the mean and variance of the Gaussian density of the emissions are used to calculate the MLEs values,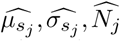 of each parameter using all the available data, since the likelihood of observing 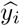 results from the contribution of each state *s*_*j*_ 

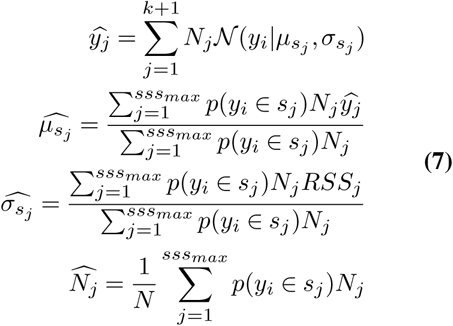

#### 3. Maximization

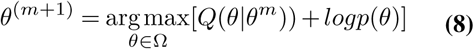

#### 4. Termination

If the difference of log-likelihood of the parameter estimate between iterations *l*[*θ*^(*m*+1)^] and *l*[*θ*^(*m*)^], exceeds a preset threshold *E*, we conclude the iterative process. 

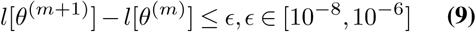

<u>step 4: MODEL SELECTION</u>

Once the parameters (mean intensity of the states, density population of the states, time locations of the transitions, and magnitude of these transitions) of the piecewise traces have been refined by the EM-clustering step, it remains to find the most likely multi-state model among all the possible states trajectories resulting from the different explored state space sizes *sss*, since we apply the MAP-EM algorithm to a state space domain that is in the range of [*sss*_*min*_, *sss*_*max*_]. While a maximum likelihood estimation approach will systematically favor the model having the most number of states because it tends to overfit the data, we preferred to exploit the same minimum description length principle used for the compression step, for the model selection final stage. The two main benefits of a MDL-based model selection approach are:

1. The trade-off between the accuracy and the model complexity, facilitating the interpretation of the model, through the parsimony principle of Occam’s razor.
2. The MDL approach is intrinsically robust to overfitting

Several model selection criteria exist, among which the Akaike Information Criterion (AIC), and its variant for small samples (AICc), and we can cite also the Bayesian version (BIC), even if it can be used to compare estimated models only when the numerical values of the dependent variables are identical for all estimates being compared, and that the BIC is only valid for sample size much larger than the number of model parameters. Moreover, even if BIC is claimed to be equivalent to MDL (57–59), we propose a corrected-MDL model selection criterion that takes into account the functional form of a multi-state model appropriately, that is, the structure of how the parameters are connected to each other, while taking advantage of the compression of the solutions domain from the two data preparation steps, according to the following objective function: 

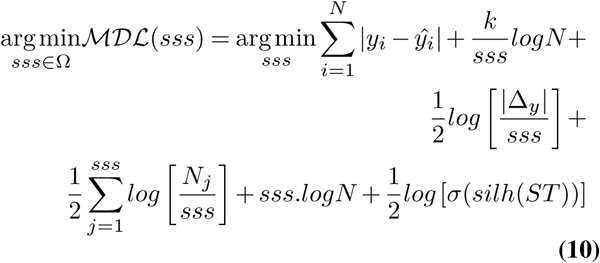

For the whole states trajectory *ST*, the silhouette coefficient *silh*(*ST*) is: 

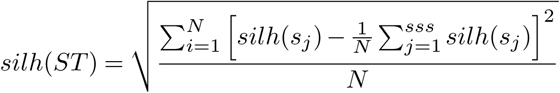

For each cluster or detected state *s*_*j*_ the silhouette coefficient *silh*(*s*_*j*_) is: 

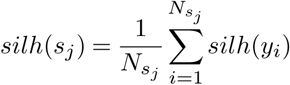

And for each single data point *y*_*i*_ the silhouette coefficient *silh*(*y*_*i*_) is: 

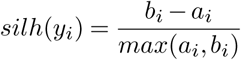

The first term in Eq. 10 represents the residuals between the compressed alone and the compressed and clustered traces, and the following terms penalize the complexity of the state-model that is compared here to a time-variant and non-linear oscillating system, by coding the description length of the oscillations rate and amplitude via the second and third terms, but also the density population of the visited states and the cost of encoding the state space size, in the fourth and fifth term respectively. Finally the last term in Eq. 10, is the silhouette coefficient *silh*(*y*_*i*_) of each observation *y*_*i*_, to measure how similar *y*_*i*_ is to all its neighbors in its allocated cluster (state) *s*_*j*_, when compared to observations in other clusters. In this intra-cluster stability metric, *a*_*i*_ is the average distance from the *i*^*th*^ point to the other points in the same cluster as *i*, and *b*_*i*_ is the minimum average distance from the *i*^*th*^ point to points in a different cluster, minimized over clusters. ℳ𝒟ℒ is optimal when the increase in the model complexity by additional states is compensated by the decrease of the residuals as measured by the goodness of fit. The added value of our model selection relies on its capability of inferring from different possible states trajectories the right state space size, even if within the sampling window, all states have not been visited at least one time, by fully integrating the stochastic complexity (60) of the reaction system.

## Results and Discussion

In this section, we investigate the robustness and reliability of the proposed cc-EM algorithm given several challenging molecular scenarios. First, we detail the performance metrics used for the validation tests and the comparison of cc-EM with other trace idealization methods. Secondly, interest is focused on the contribution of each stage of cc-EM and the added value of their combination into the resulting learning algorithm for the optimization of the trace idealization. Then, complexity is added at each experiment level by scanning the signal parameters space wrt the molecular kinetics profile and the sensor’s characteristics. Finally, we resume the performance profiles of cc-EM and propose in Sup. Note 4 (simoVIS: a SIngle-MOlecule kinetics VISualization Tool) an alternative single-molecule data visualization tool to explore the underlying transition-state network of the modeled smFET signals.

### D. Performance metrics

The table in Sup. Note 1 resumes the Performance Metrics used to assess the robustness of cc-EM and to compare it to other single-molecule trace idealization methods, among which the RMSE between the estimated idealized trace and the ground truth, the macro-averaged precision, recall and F-score (61–63). We also propose an original hybrid metric, the OKB score, as the **O**verall **K**inetic **B**ias of the resulting state-model parameters inferred from the obtained idealized trace.

### E. The combination compression - clustering - EM refinement optimizes significantly the smFET trace idealization

We compared the goodness of fit through the RMSE metric between 200 synthetic states trajectories of 2500 data points each and the estimated idealized trace obtained at each stage of cc-EM, given different noise levels (figure 4a) and a varying model complexity (number of hidden states) (figure 4b). While the outputs obtained by the MDL-based compression alone are comparable to those of the compressed and clustered (MDL+k-medoids) step for relatively medium and low noise levels (*SNR ≥* 5*dB*), the MDL+k-medoids outperforms the MDL algorithm in (40) by two orders of magnitude in terms of accuracy for highly noisy environments (*SNR* = 1*dB*). Concerning the robustness of cc-EM to the number of hidden states to detect, each additive step of the proposed algorithm brings closer to the best reachable output that we can get from a HMM trained with the true initial priors (true transition probability matrix) of the simulated traces, by significantly reducing the residuals between the estimated idealized trace and the ground truth. Finally, whatever the noise level and the molecular model complexity, the combination compression+clustering+EM refinement always leads to more accurate idealized traces than each separate stage of the hybrid cc-EM algorithm.

**Fig. 4.**
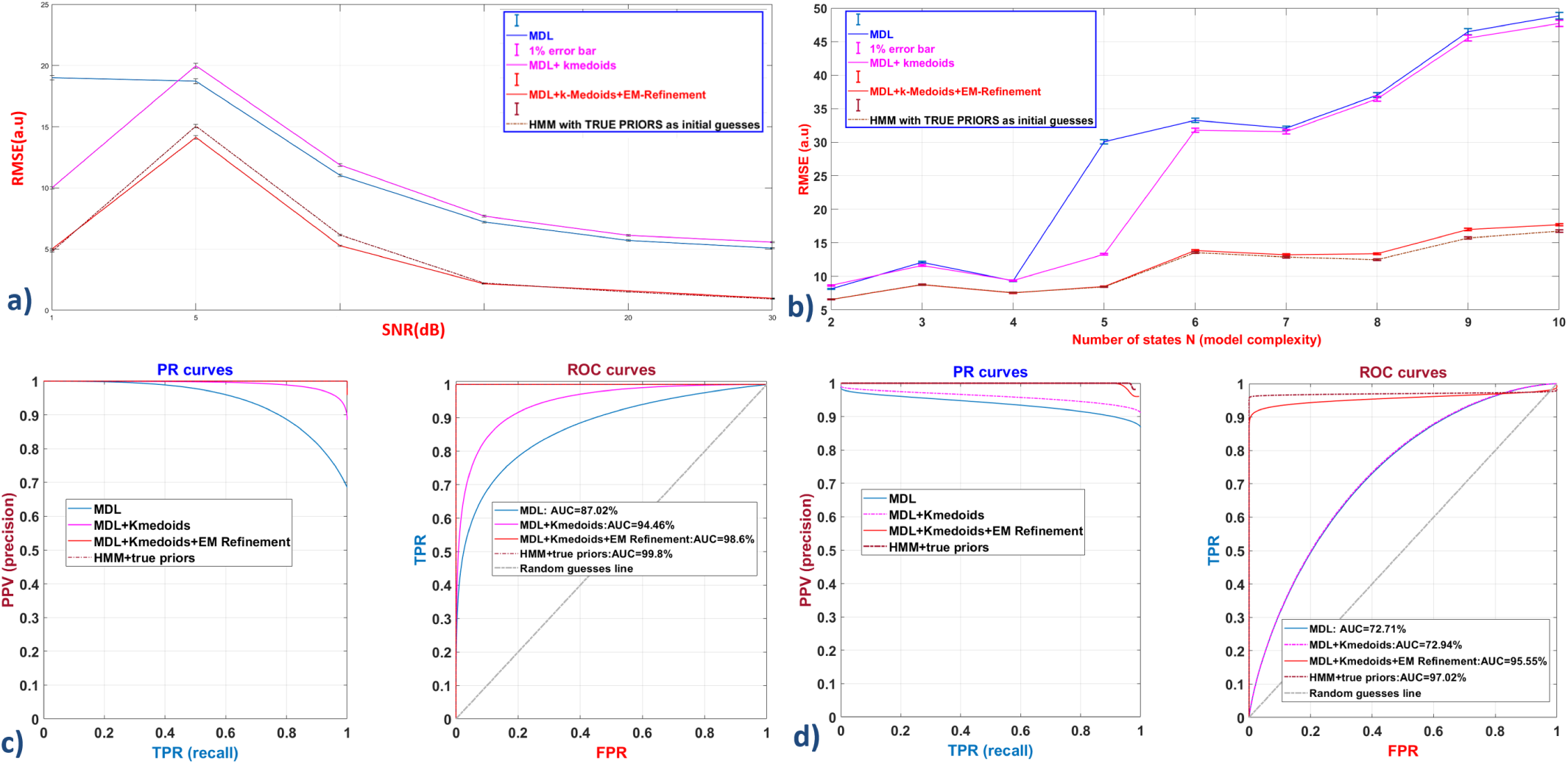
Performance comparison between each additive step (compression-clustering-EM refinement) of the proposed cc-EM trace idealization method, and the HMM outputs obtained with the true priors as initial guesses.a)Goodness of fit comparison of each step of the cc-EM algorithm given a varying noise level (1 *≤ SNR ≤* 30*dB*) corrupting a 2-states first-order Markov process. The best reachable RMSE score are given by the HMM outputs trained on the true simulated transition matrix. b) RMSE comparison of each step of cc-EM given an increasing model complexity (2 *≤ sss ≤* 10) and a constant noise level of 1dB. c)ROC curves for each step of cc-EM given a varying noise level (1 *≤ SNR ≤* 30*dB*) on a 2-states first-order Markov process left: Precision-Recall curve, Right: True positive rate (TPR) vs False positive rate (FPR) curve. d) ROC curves for each step of cc-EM given an increasing state space size (*sss*) (2 *≤ sss ≤* 10) and a constant noise level of 1dB. left: Precision-Recall curve, Right: True positive rate (TPR) vs False positive rate (FPR) curve.

Secondly,since the single-molecule trace idealization can be seen as both a stepwise regression task or a classification/clustering of the signal data points into a known or learned number of classes or clusters, each corresponding to a molecular state, that is a plateau in the state trajectory, we assessed the performances of our method in terms of precision and recall scores, and via the AUC of the ROC curves in figure 4. For each tested method, the resulting ROC curves have been calculated from the average true positive and false positive event detection rates, via the corresponding confusion matrices (64) obtained from the multi-class classification of the detected states in 200 synthetic traces of 2500 data points each, given different noise levels (figure 4c) or an increasing number of hidden states (figure 4d). While the AUC scores are comparable for the MDL-based compression (72,71%) and the MDL+k-medoids step (72,94%) as the model complexity increases (2 *≤ sss ≤* 10), the latter is clearly a better classification model for different noise levels (1 *≤ SNR ≤* 30*dB*), with an AUC of 94,4%, than the MDL-based clustering approach (AUC=87%). Again, the combination of the three steps compression+clustering+EM refinement systematically leads to more and highly accurate classifiers with an AUC of 98,6% and 97% under an increasing noise level and model complexity respectively.

### F. Robustness to the sensor profile

Figure 5 resumes the RMSE boxplots between the learned state trajectories and the simulated ones, given different tested sensor parameters and molecular kinetic profiles.

**Fig. 5.**
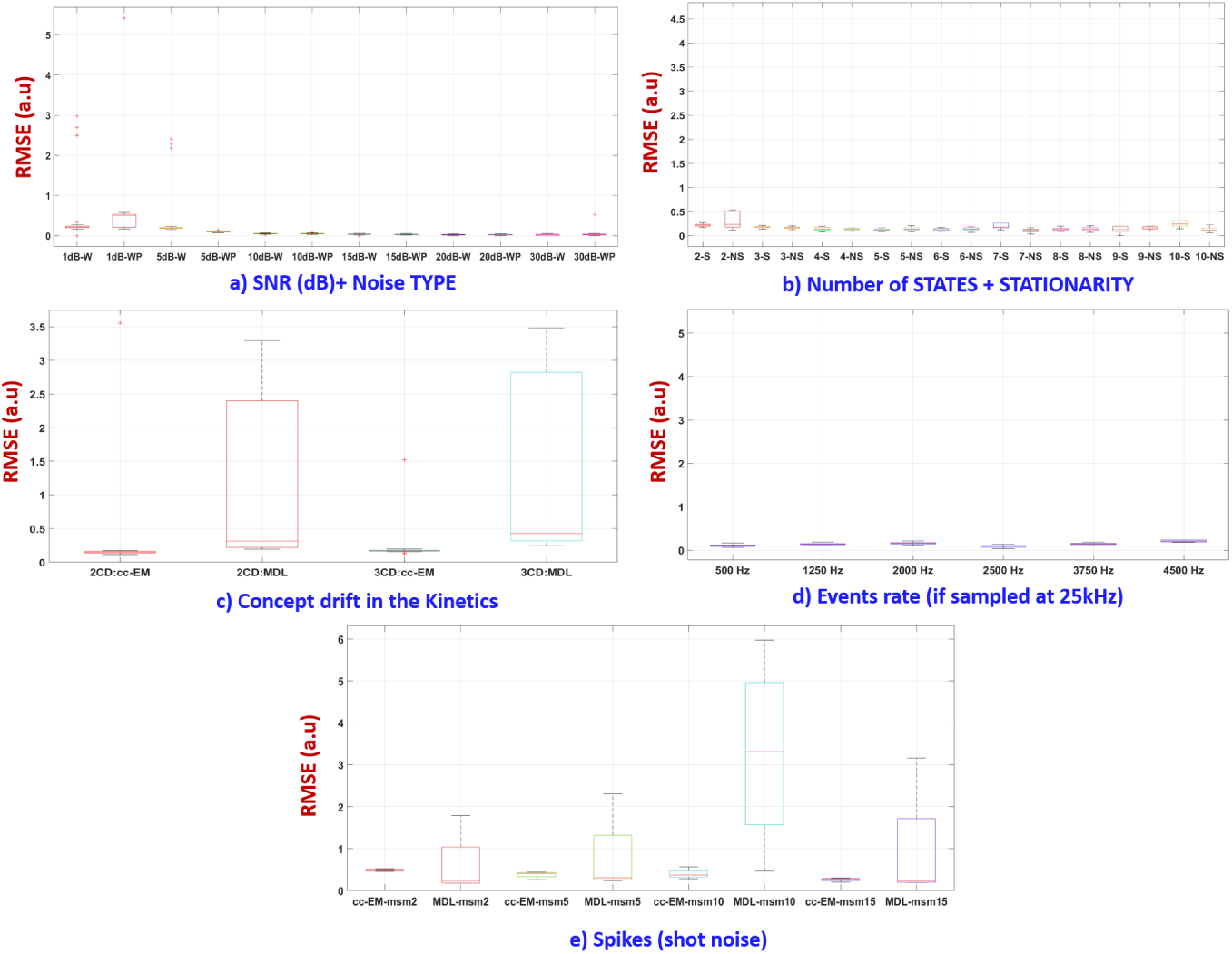
Robustness of the cc-EM idealization algorithm for various signal and kinetic parameters: a) intensity of Gaussian noise (W) and mixed to pink noise (WP), b) number of stationary (S) and non-stationary (NS) sources, c) concept drifts in 2-state transition rates (ex. 2CD-cc-EM for transitions between kinetics of 500 and 1250 events tested with our cc-EM method, versus the MDL alone proposed in (40), while 3CD-cc-EM denotes transitions between kinetics of 500,1250 and 2500 events) d) robustness of cc-EM under fast kinetics e) Robustness of cc-EM to false detection under random spikes from impulse noise

#### F.1. Robustness to the Noise Level and Mixed Noises

To assess the robustness of cc-EM to the noise level we tested it on 360 synthetic traces of 2500 points each, evenly distributed according to the SNR value and the type of noise. figure 5 a), shows how cc-EM is robust and accurate even under highly noisy environments (SNR=1db), while in figure 4a, one can better visualize the effect of each step of the proposed cc-EM algorithm, and how well the data preparation steps by the MDL-based compression and the K-medoids clustering enable to reduce the RMSE significantly under high noise levels (SNR=1db), making the cc-EM algorithm more efficient than the different steps (Baum-Welch and Viterbi) of conventional HMM even if trained with the true transition matrix as priors. smFET traces corrupted by mixed noises (AWGN and flicker) exhibit locally some changes of their statistical properties (mean intensity and variance) in a manner that can complicate the detection of the states trajectory, leading to wrong state-model parameters’ values (average conductance of the state levels, population density of states, position and amplitude of the transitions) or worse to an erroneous molecular model (under/overfitting of the number of hidden states). For this reason, we tested the performances of the proposed cc-EM algorithm under mixed noises (AWGN+flicker), taking into account the signal distortion due to the measurement noise from the acquisition chain, but also due to the process noise from the kinetics of the molecular dynamics under monitoring. As a result, figure 5 a) shows the robustness and accuracy of cc-EM for trace idealization under such varying noise variance.

#### F.2. Robustness to the Trace Length

The learning rate of cc-EM depends on finding an optimal signal compression ratio to reach the best piecewise regression fit of the raw signal, while the prediction rate of the right state space model corresponding to the right number of hidden states depends on the sample size that must have a minimum threshold number of data points to correctly infer the molecular model from the learning dataset. The effect of the sample size parameter on the learning performances is also correlated to the involved kinetics of the molecular dynamics, namely the transitions rate between states, so that the more frequent are these transitions the “richer” will be the informative content of the resulting states trajectory, even if all the states have not been visited at least one time within the sampling window (imbalanced multi-class classification of the states). Our proposed cc-EM algorithm needs a training dataset of at least 2500 points to reach optimal accuracy scores as detailed in the performance landscapes in figure 7 for fast kinetics (1 event every 3 data points) and noisy environments (*SNR ≤* 1*dB*).

**Fig. 6.**
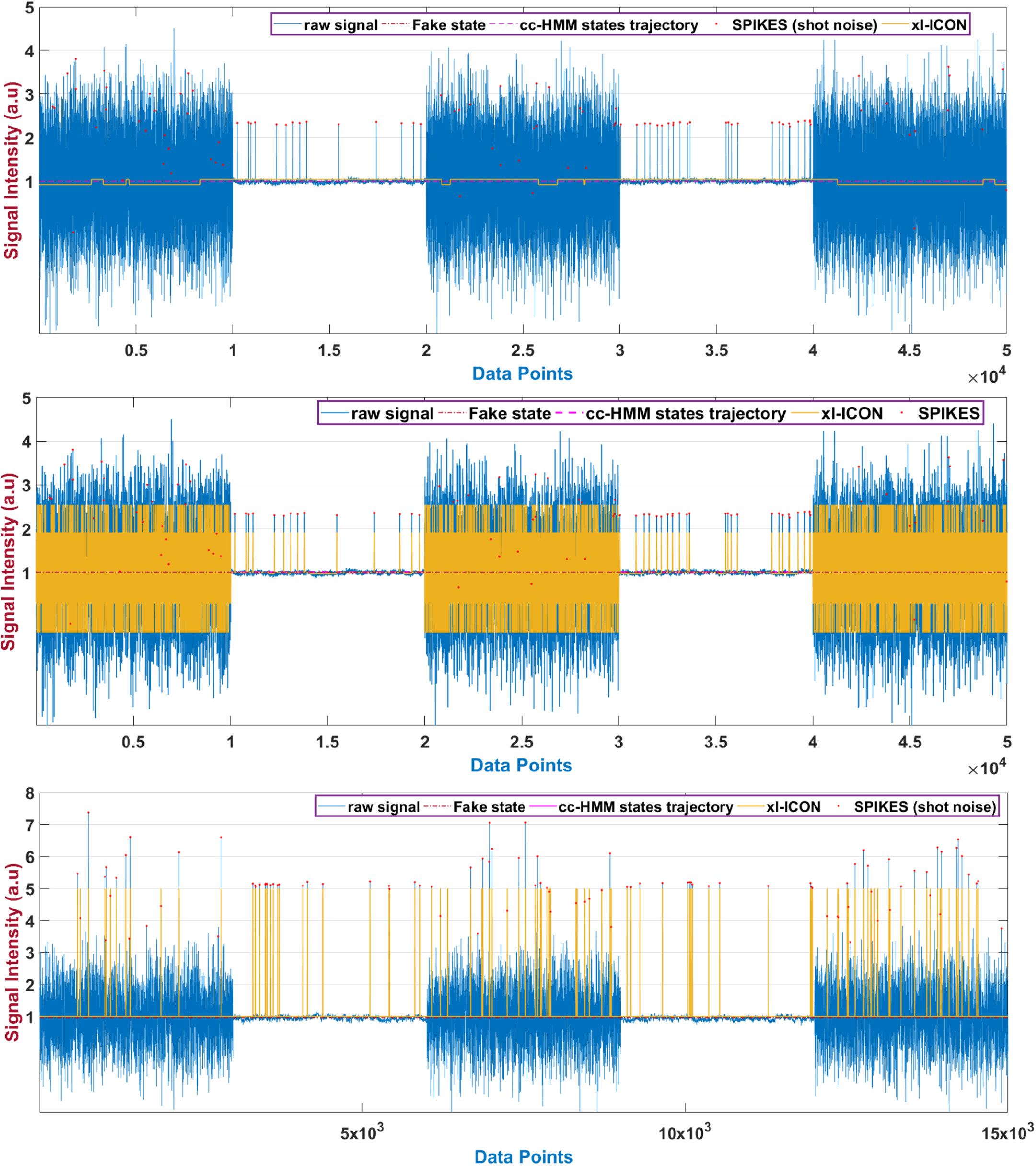
Robustness comparison to shot noise between cc-EM and a BNP algorithm. A synthetic single state has been corrupted by AWGN, flicker noises given a SNR=1dB, and 100 spikes randomly placed along the time series to check the specificity of the proposed cc-EM algorithm and its robustness to false positive events and states. Top: cc-EM managed to retrieve the single fake state while the BNP method with 500 MCMC iterations led to a 2-states model. Middle: As we increase the number of MCMC iterations, the BNP algorithm overfits more, while the final estimated number of states is 8 for 5000 MCMC iterations in this case. Bottom panel: In this other challenging synthetic trace, we increased the average amplitude of the spikes by two orders of magnitude related to the mean intensity of the raw signal corrupted by mixed white Gaussian and flicker noises, given a SNR of 1dB. Again the BNP systematically overfits the model complexity (3 hidden states detected instead of 1) by assimilating the spikes as being transitions towards short-lived states, while cc-EM remains robust thanks to its bit-length encoding scheme of the state model parameters that doe not only depends on a the signal level dimension

**Fig. 7.**
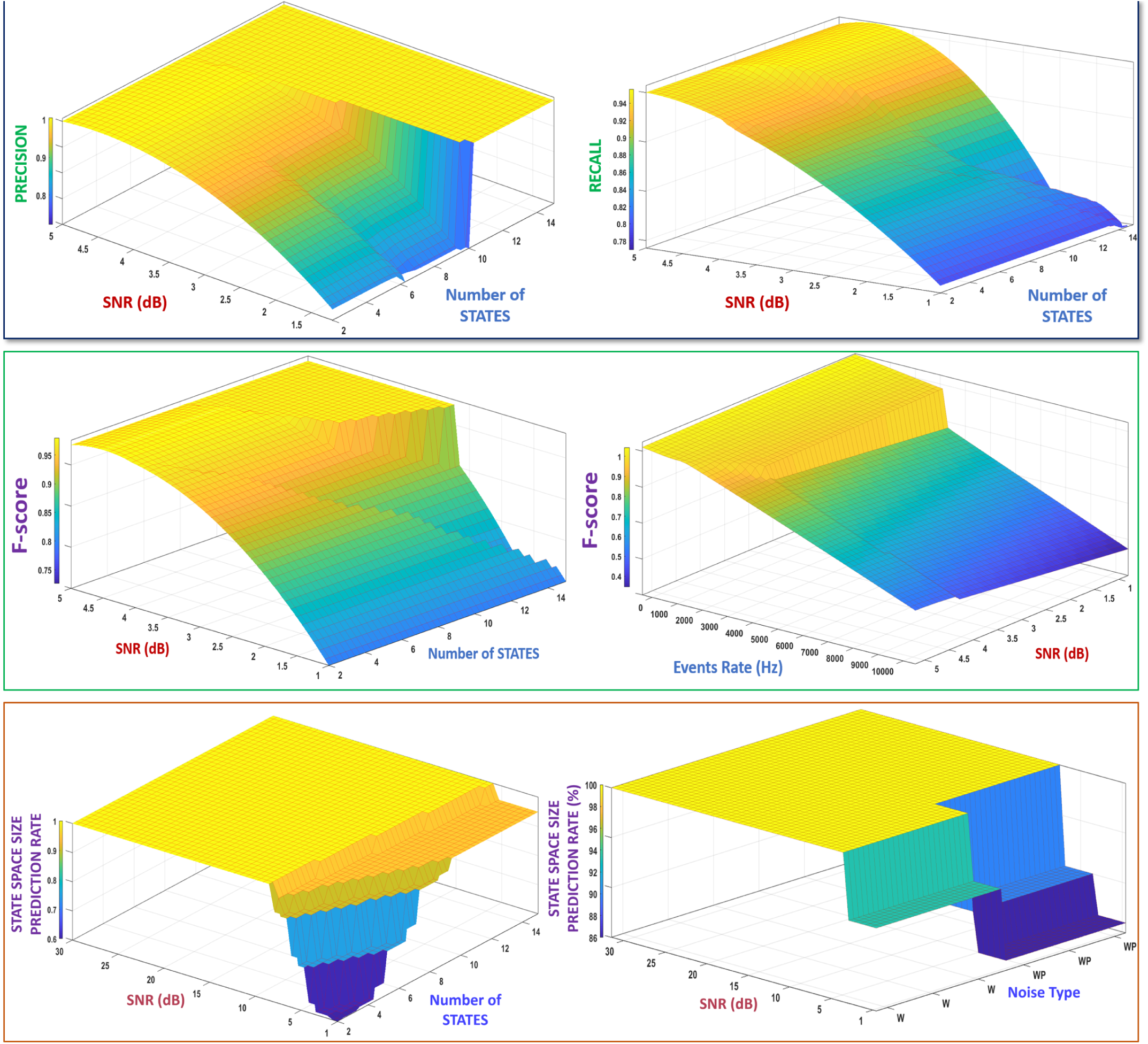
Performance profile of the cc-EM algorithm through 3D landscapes of its application domain that guarantees the most accurate outputs given the sensing and kinetic parameters. Top left panel:precision of cc-EM wrt the noise level and the kinetic model complexity. Top right panel: recall of cc-EM given a varying noise level and model complexity. Middle left panel: F-score given different noise levels and number of hidden states. Middle right panel: F-score of cc-EM given the noise level and the event to measurement ratio that is in the ranges of [1*/*8, 1*/*3], corresponding to one molecular event every 8 or 3 data points in the recorded signal. Bottom left panel: prediction rate score of the MDL-based model selection step in cc-EM, given different noise levels and model complexity. Bottom right panel: prediction rate of the same model selection for different noise types and levels.

#### F.3. Robustness to the Sensor Baseline Drift

Because of the stochastic nature of the sensed molecular dynamics, long acquisition periods and high-throughput measurements are necessary to record as much data as possible for statistical analysis, and to collect enough training samples for the cc-EM learning algorithm. Due to such sampling conditions, the sensor baseline may drift significantly during the acquisition period, and potentially induces fake states and transitions in the smFET time trajectory, leading to wrong kinetic estimates. Our proposed framework includes a baseline drift compensation tool (41) to remove such parasitic signal from the useful signal corresponding to the sensor response to the probed molecule. Based on Information Theory, this model-free baseline wander removal method consists of an iterative multiscale signal compression technique implemented into a blind source separation scheme, to decompose the multi-component smFET signal into several embedded signal layers, among which is the sensor baseline. The resulting baseline drift compensation is tailored to separate accurately continuous and discrete time-variant signals from each other, given a robustness to high and mixed noises, concept drifts, and various shapes and rates of baseline drifts, even if the variance of the noise and the baseline are different, and with-out prior knowledge on the sensor parameters or the molecular kinetics. To better visualize the effect of the baseline drift on the trace idealization performances,see figure 12 in Sup. Note 3 (Trace Idealization under Baseline Drift).

**Fig. 8.**
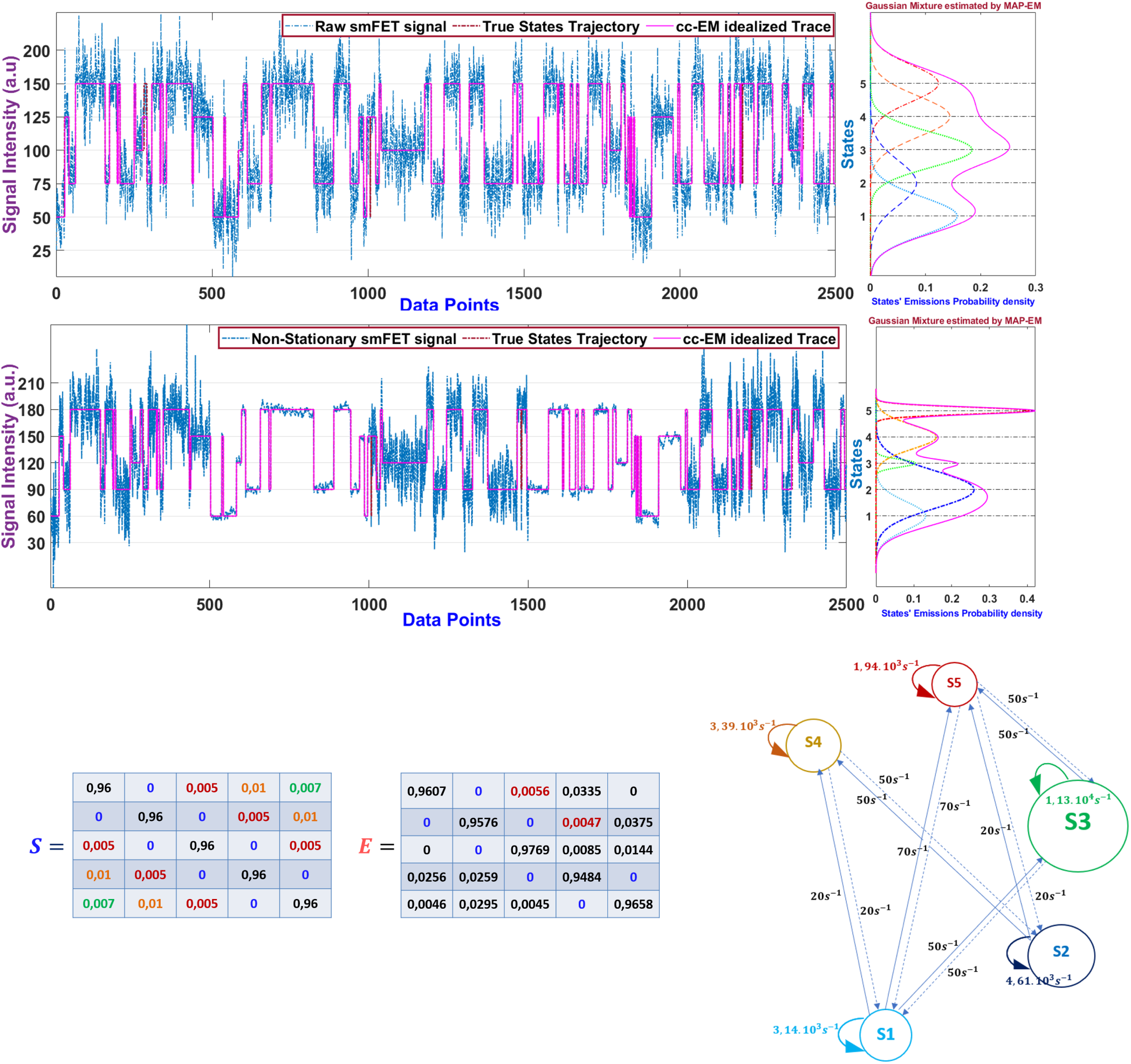
a 5-states model with a Toeplitz transition matrix, and the calculated rate given a sampling frequency of 25kHz, obtained by the simoVIS single-molecule data visualization tool. *Top left:* Comparison between the simulated state trajectory (dashed red) and the cc-EM idealized trace (pink). *Top right:* The corresponding emission distributions of each state and of their time trajectory (pink), obtained by the MAP-EM step of cc-EM (i.e. initialized with the state sequence obtained from the compression and clustering stages). The dotted horizontal lines indicate the estimated mean values of the emission distributions. *Middle Panel:* The same states trajectory has been corrupted by a mixed AWGN+flicker noise given a SNR of 1dB, and we compared the idealized trace obtained by cc-EM to the simulated ground truth. *Bottom left:* Comparison between the estimated transition matrix (E) and the simulated one (S). *Top right:* Diagram of the transition rates highlighting some forbidden transitions, while each sphere correspond to a state whose radius is proportional to its density population.

**Fig. 9.**
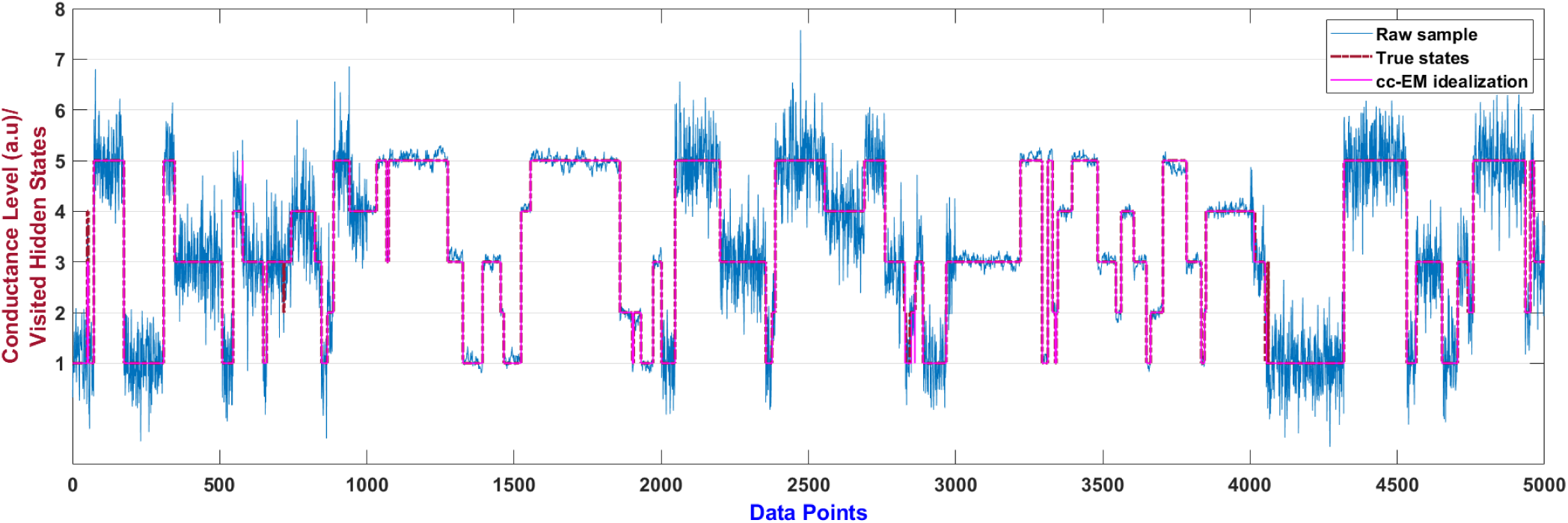
A simulated 5-states model (red) corrupted by a mixed (AWGN+Pink) noises (SNR=1db) (blue) with the estimated states trajectory (magenta) obtained by the cc-EM algorithm

**Fig. 10.**
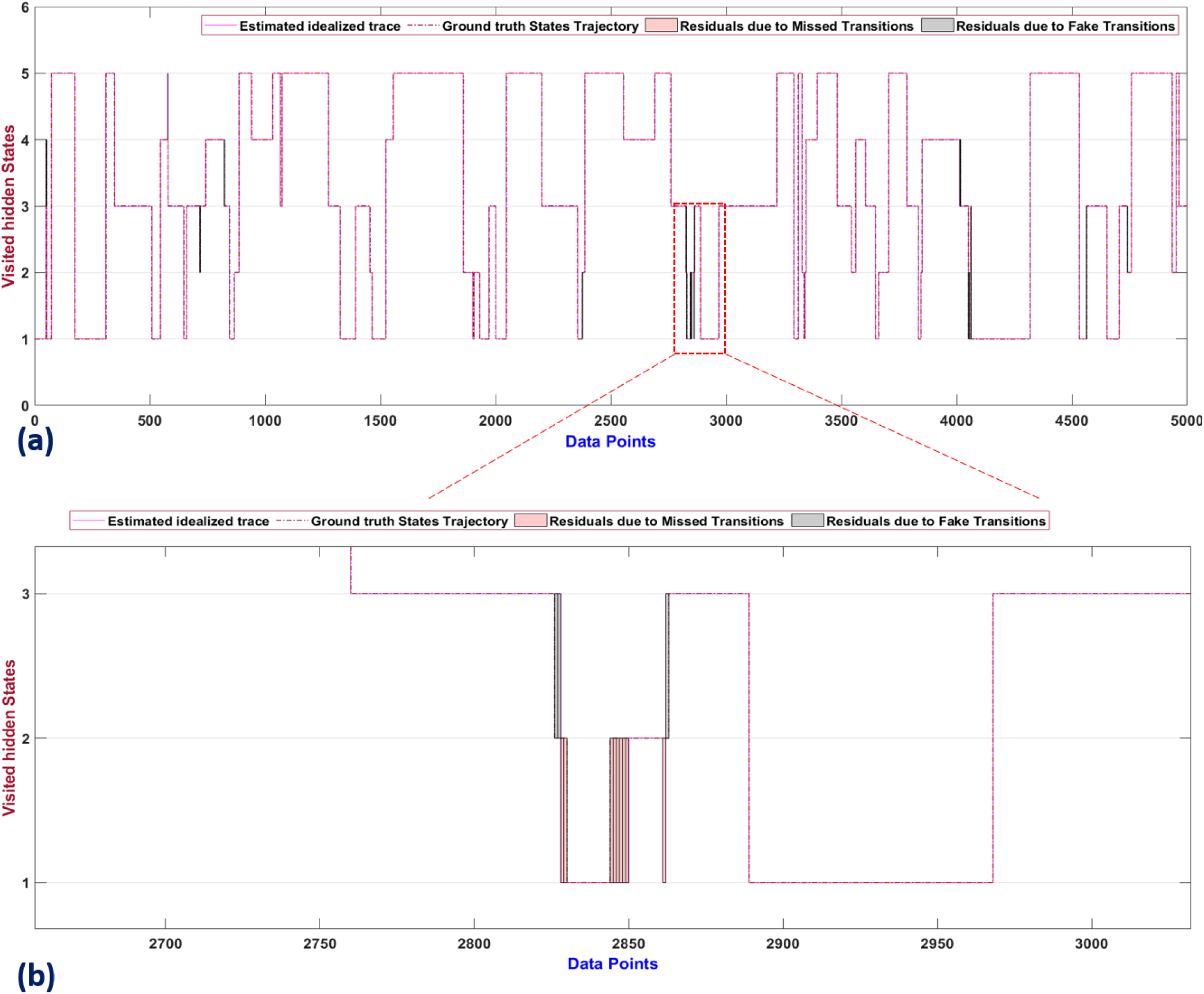
(a):Comparison between the estimated idealized trace and the ground truth states trajectory in terms of cumulative integral of the residual areas generated by all the missed and fake transitions (b) zoom-in section on each of the false positive (grey shaded areas) and false-negative transitions (pink shaded areas) with a single point time resolution.

**Fig. 11.**
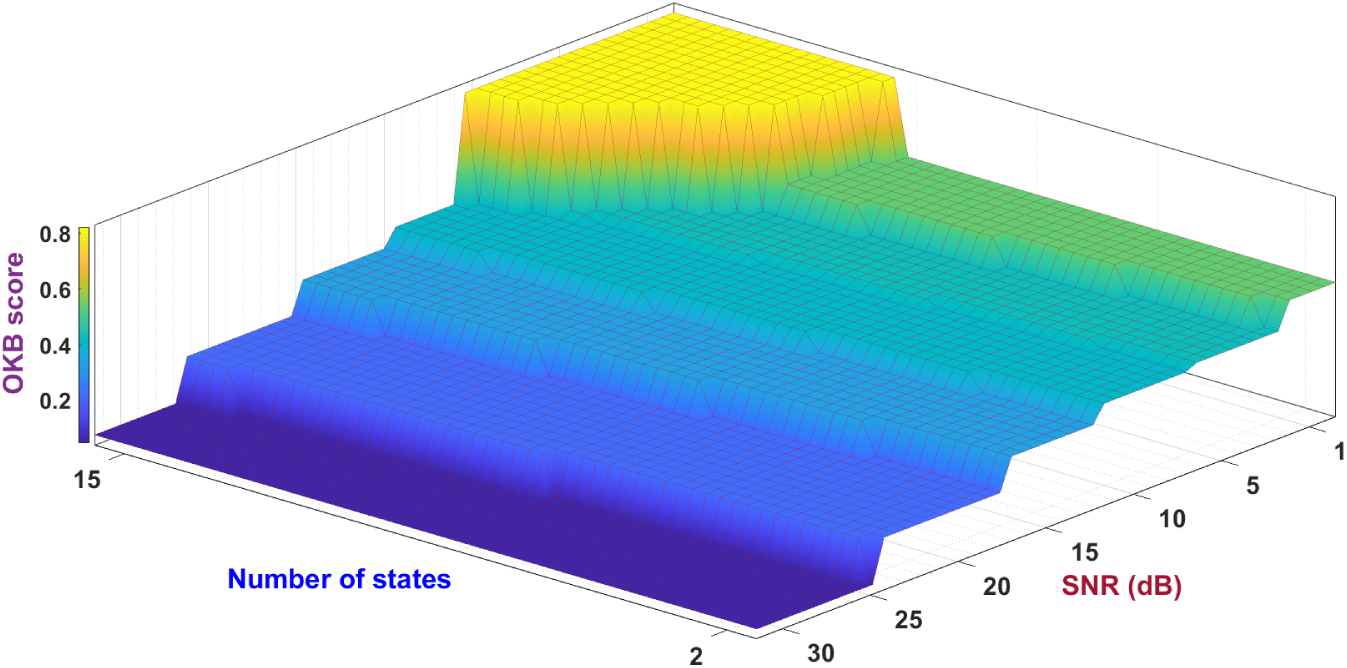
OKB score landscape of the cc-EM algorithm to assess its performances in terms of overall accuracy of the idealized trace, the kinetic parameters estimates and the prediction rate of the model selection step to infer the right molecular state space size, given varying model complexity and noise level.

**Fig. 12.**
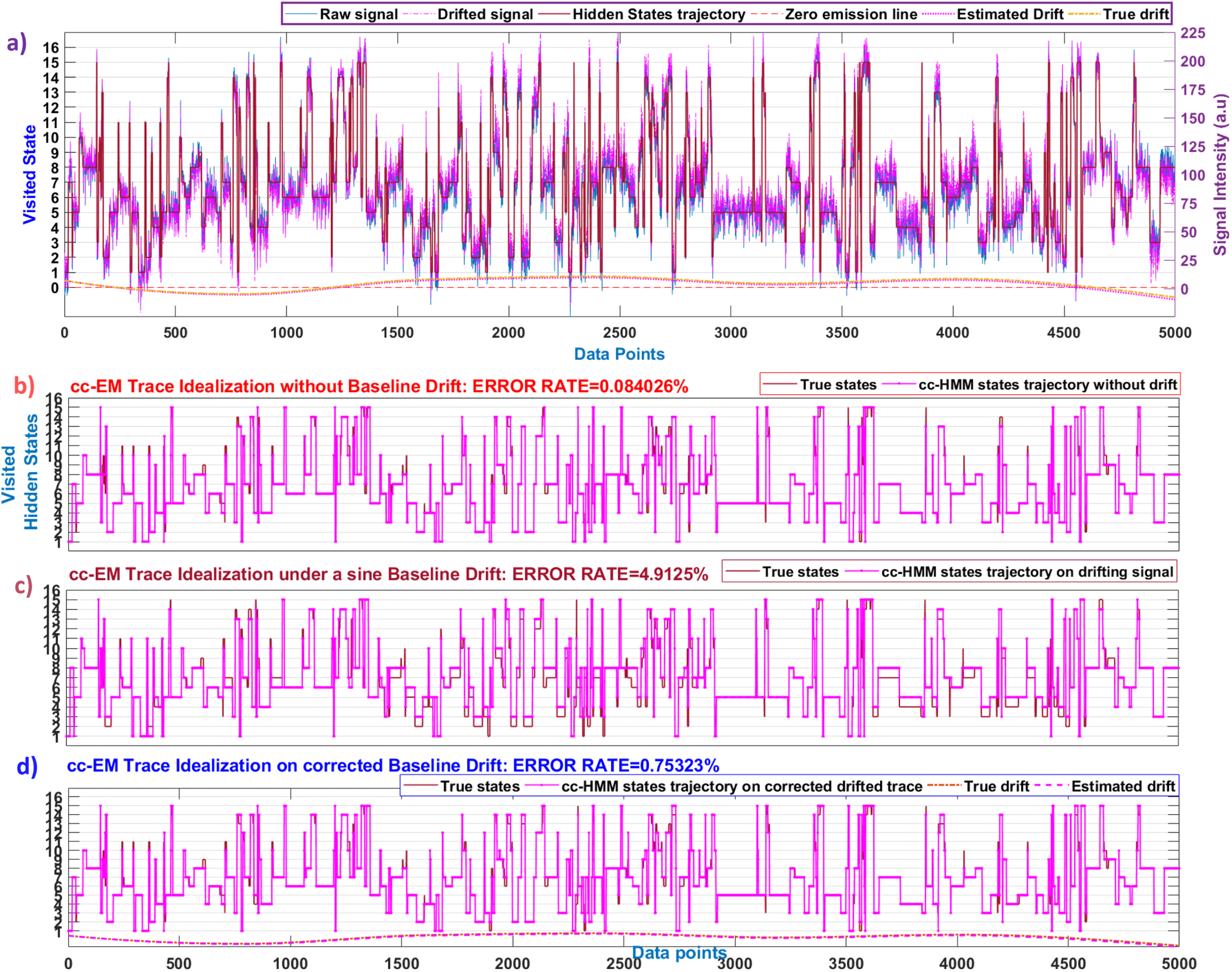
Trace idealization of a simulated 15 states-model under a sine sensor baseline drift that locally inserts fake steps and transitions in the resulting states trajectory. a) smFET trace corrupted by an AWGN with a SNR of 1dB, and a sine drift, leading to the drifted source signal (pink). The true states trajectory and the simulated baseline drift are also depicted. b) comparison between the cc-EM idealized trace and the ground truth, without any baseline drift, leading to an error rate (percentage of missed transitions in time and/or amplitude related to the total number of true transitions) of 0.1% c) the same comparison under a baseline drift that increases the error rate of around 5% despite a very smooth sine drift with with much smaller oscillations amplitude compared to the local states’ noise variance. d) the same comparison after a baseline drift compensation that significantly limits the impact of the parasitic signal on the cc-EM performances.

#### F.4. Robustness to shot noise and outliers

We also tested the specificity of cc-EM through the false positive detection rate of short-lived states that can be confused with the impulse noise. For each simulated trace we inserted 100 impulses with varying amplitude (between 5 and 10 fold the local noise variance at the time location of the shot noise), and randomly placed all along the trace given different model complexity (2,5,10,15 states), while all the resulting traces have been corrupted by a mixed noise (AWGN+pink) with a SNR=1dB. We calculated the distribution of the RMSE between the estimated state trajectory and the ground truth, and the prediction rate of the state space size to check how cc-EM is robust to overfitting. We found that cc-EM correctly predicts the number of states in 91% of cases, given a higher robustness to false positive detection than the MDL algorithm in (40), and with much less estimate errors as can be seen with the RMSE boxplots 5. We also compared our proposed cc-EM algorithm to a Bayesian non parametric approach using the xl-ICON framework (65) to assess their specificity wrt false positive event detection and their robustness to overfitting under mixed noises (AWGN+pink+shot). To do so, we simulated a single fake state with 100 randomly positioned impulses simulating fake events, and sequentially corrupted the resulting trace by both AWGN and pink noises, given a SNR of 1dB, which emphasizes the magnitude of the transitions by the time varying noise variance, and thus increasing the risk of detecting false positive short-lived states. Figure 6 clearly shows that the cc-EM algorithm exhibits a higher specificity against fake transitions and states, leading to more accurate estimates of the state space size and the states trajectory, while the BNP method tends to overfit systematically as we increase the number of MCMC iterations, which is a user-preset parameter not learned from the data, making such approach intrinsically semi-supervised, but especially more sensitive to shot noise than the cc-EM method. Such robustness to outliers and high specificity of the proposed cc-EM algorithm relies on the encoding scheme of the state model parameters, instead of defining the states and their transitions from each other only wrt the signal magnitude dimension. In fact, thanks to the information theory approach, we manage to learn the intrinsic dimensionality of the smFET time series (66), while most of the existing trace idealization methods focus on the signal levels and/or on some distributions associated to the emissions of the hidden states.

### G. Robustness to the molecular kinetics profile

#### G.1. Robustness to the Event Rate and Fast Kinetics

To assess the robustness of cc-EM to fast kinetics corresponding to the frequency of the transitions between the molecular conformational states, wrt to a sampling rate of 25kHz in our simulations, we tested several event rates, by modulating the transition probabilities between a 2-states first-order Markov process, whose the resulting state trajectory has been corrupted by a mixed noise (AWGN+pink,SNR=5db). Each boxplot in figure 5d represents the distribution of the RMSE between the estimated state trajectories and the simulated one, for 30 traces of 2500 points each, while the states’ dwellings are not equally probable, which increases the difficulty level of the trace idealization objective. On average, the resulting density population of states from such fast kinetics, is so that the number of data point available to define a state given a sample size, is about 1% of the total trace length, without assuming that within the sampling window, all the simulated states have been visited at least one time (imbalanced multi-class classification of the states ie one or more states have very low proportions in the training data as compared to the other states). Even under a time-varying noise variance that complicates the states parameters estimation (especially the mean intensity of each plateau in the idealized trace), our cc-EM method remains highly accurate and reproducible for the detection of fast molecular dynamics (up to 1 event every three data points), within the right state space size domain.

#### G.2. Robustness to the State Space Size

To assess the reliability of cc-EM to correctly estimate the states and rates parameters given different number of hidden states, we calculated the goodness of fit through the RMSE (figure 4 b) for each step (compression, clustering, inference) of the cc-EM algorithm, and compared the resulting RMSE values with those obtained by a HMM trained on the respective true transition matrix of each simulated trace as priors. In figure 4 d), the precision-recall and the ROC curves have been calculated given an increasing model complexity. We also calculate the prediction rate of the model selection step given an increasing number of hidden states and different designs of transition matrices (Toeplitz or asymmetrical matrices). For instance, for a 15-states model, the F-score is about 0.95 for a SNR=1dB and the prediction rate of the right state space size is about 92% for white Gaussian noise and 86% in the case of mixed noises. All these validation tests demonstrate how cc-EM can handle complex single-molecule time trajectories, while its computational time (*𝒪*(*N* log *N*)) makes it applicable to large dataset whatever the model complexity, unlike the forward-filtering backward sampling algorithm whose complexity relies on both the sample size and the number of states (*𝒪*(*NK*^2^)).

#### G.3. Robustness to concept drifts and non-stationarity

Because of the possible non-stationarity of the smFET recordings, it is important to assess the accuracy of cc-EM under concept drifts corresponding to changes in the data generating process itself, namely at the kinetic level of the probed molecular dynamics. To this end, we simulated a 2-states random walk whose the transition matrix evolves over time resulting in a state trajectory having either 2 concept drifts making the reaction system sequentially swap between 500 and 1250 events per second for a sampling rate of 25kHz, or 3 concept drifts between kinetics of 500,1250 and 2500 events given the same sampling conditions.In figure 5 c), we plotted the boxplots of the RMSE between the estimated state trajectories and the corresponding ground truth, for 60 traces of 2500 data points, and compared cc-EM to the MLD-based method in (40). For each concept drift scenario, cc-EM leads to better robustness and accuracy than the MDL alone outputs. In 5 b), we tested the accuracy of cc-EM related to the number of molecular states and compared the RMSE between stationary and non-stationary sources, and found that cc-EM remains robust and highly accurate whatever the types of sources. Finally, the detection of concept drifts along the time series and the automatic update of the state-model parameters accordingly, beyond the scope of the present study, will be addressed as a future work, while the resulting idealized traces and their corresponding state space size, obtained by cc-EM under such non-stationary environments, remain much more accurate than conventional hidden Markov model-based methods and variational Bayesian inference.

### H. Performance profile of the cc-EM algorithm

In this section we resume the performance profile of the cc-EM algorithm into 3D charts enabling to better visualize the landscape of the application domain of cc-EM that leads to the best accurate outputs wrt the Performance Metrics for the trace idealization and kinetic parameters estimates. Figure 7 depicts in the top panel the precision and recall landscapes of cc-EM, given varying sensor and molecular kinetics parameters. The precision score of the built-in event detection method in cc-EM remains constant and high (*<* 5% of false positive rate) for a SNR of 5 dB and lower, that are typical noise levels observed in smFET experiments. Surprisingly, beyond a threshold number of 10 states, our changepoint detector becomes insensitive to the noise level and the model complexity (for a time-constant and a relatively low event rate between 4 −8% of the trace length), so that no decaying effect on the precision score can be observed any more, as the false positive rate vanishes, as if the more oscillating is the signal due its number of states, the better the detector learns from these more frequently changing patterns (the shape and structure of the states trajectory), and manages to reduce the false positive detection rate. Concerning the recall score landscape given both the noise level and the complexity of the molecular dynamics, our idealization method presents a high sensitivity (*≥* 95% of true positive rate) for a SNR of 5dB and more, and decreases when the SNR tends to 1dB with a score of 75%. We do not see the same counter effect on the recall score as for the precision score, when the number of states increases, indeed the sensitivity remains unchanged for a SNR=1dB whatever the state space size.

The middle panel of figure 7 resumes the trade-off between specificity and sensitivity through the F-score, which is the harmonic average of the precision and recall, and represents the accuracy of the trace idealization method, given the noise level and the number of molecular states. The accuracy of our proposed method remains high (≥ 98%) and stable for a moderate noise while it progressively decreases under very noisy environments (*SNR ≤* 1*dB*), but with a quite acceptable value (80% of accuracy), whatever the state space size. In the right panel, the F-score landscape is depicted given the noise level and the event rate (for a sampling frequency of 25kHz). Up to 3000 events per second, (1 event every 8 data points for a sampling rate of 25kHz), the accuracy remains above 90% even under a SNR of 1dB, while in the extreme case of 8000 events per second (1 event every 3 data points for a sampling rate of 25kHz), the accuracy of cc-EM is about 65% if the noise level is high (SNR=1dB) and is equals to 70% if SNR=5dB.

The bottom panel of figure 7 represents the prediction rate of the model selection algorithm to successfully estimate the correct number of hidden states given both the noise level and the model complexity of the simulated molecular dynamics (left panel), and given the SNR value and the noise type, AWGN or mixed AWGN+flicker noises (right panel). The 3D chart depicts the behavior of cc-EM along the signal and molecular kinetics parameters space, and shows its robustness and accuracy for a SNR of 5dB and higher with a prediction rate of 90%, and between 60% and 80% for a SNR of 1dB. We retrieve the same behavior to the noise level beyond the threshold number of 10 hidden states, as explained for the precision score landscape in the top panel of 7, so that the more states to detect the more reliable the model selection is, even under high noise levels (1dB) because the encoding of the event rate in the cost function of the model selection algorithm as an oscillation parameter that is correlated to the state space size through the transition matrix, helps the model selection algorithm to better converge to the right molecular state space.

## Application to drifting smFET signals

To benchmark our proposed cc-EM algorithm, we apply it to the analysis of synthetic smFET data, using a random walk through a Toeplitz transition matrix, and corrupted the resulting trace first by an additive white Gaussian noise (AWGN) (Fig.8 upper left panel), and then by a mixed AWGN+flicker noise given a SNR=1dB (middle panel). Not only cc-EM recovered the hidden states sequence with only a very few missed transitions (4 false negative in the case of AWGN, and 3 missed events under a time-varying noise variance), but also without any false positive states and transitions, while the learned transition matrix (E matrix in the lower panel of figure 8) has a relative low bias compared to its true value (S matrix).If we compare the states’ emission distributions between the two smFET signals, we notice the significant overlap across the non-stationary emissions spectra that complicates the states identification. As previously mentioned in the section Robustness to the Event Rate and Fast Kinetics, cc-EM can handle imbalanced multi-class classification of states that are so rarely visited within the sampling window (about 1% of the total trace length) that they could be ignored with conventional HMM-based trace idealization methods.

## Conclusions

cc-EM is a complete single-molecule data analysis framework tailored for non-stationary and drifting smFET trace idealization, and the kinetics modeling of the biomolecular dynamics between conformational states. This alternative entropy-based piecewise segmentation process highlights the preponderant electrical signatures corresponding to the dynamics of the probed biomolecules, even under low SNR and mixed noises, including AWGN, flicker and shot noises.

More importantly, the proposed cc-EM algorithm can learn the states and rates of Markovian processes without detailed balance (asymmetrical transition matrices), but also of non-Markovian chains, while the built-in model selection can infer the right state space size of the corresponding molecular model even in the case of imbalanced states sparsely populated, and under some particular noise and kinetics conditions, cc-EM achieves the same performances even if within the sampling window, all the states have not been visited at least one time.

Since cc-EM is a model-free framework it is compatible with reactions that do not follow a particular kinetic scheme, and because cc-EM does not rely on any assumptions for the states and rates characterization, it is applicable whether the system is in or out equilibrium, taking advantage from the time resolved smFET sensors’ measurements to provide the individual transition paths of a molecule’s trajectory.

We implemented our algorithm in a Matlab framework with several companion tools among which, a baseline drift removal plugin, and a single-molecule FET trace generator with options to setup signal and molecular kinetics’ parameters. Option to store all the idealized and corrected baseline traces with their statistics is also provided.

Validation tests on a wide single-molecule signal parameters space conclude to the robustness and reliability of cc-EM to precisely and accurately learn the states and rates from the sensed molecular dynamics, while the proposed trace idealization method can be generalized to other single-molecule sensing techniques (smFRET, patch clamp recordings) without prior knowledge on the data generating process nor signal prefiltering.

## Software Availibility

cc-EM can be downloaded and installed automatically as a Matlab desktop application or a standalone executable by following the instructions in the user guide (code and signals available on request).

## Supplementary Note 1: Performance Metrics

## Supplementary Note 2: OKB score: a new overall accuracy metric for the performance evaluation of single molecule trace idealization

Beside the precision and recall metrics to assess the sensitivity and specificity of a binary or multi-class classifier, as the trace idealization algorithm is and does to allocate each data point of the raw signal to a learned or a priori defined number of states, we can also evaluate how well or not the transitions have been estimated in term of time location, amplitude, and number. To do so, we propose the OKB metric to calculate the Overall Accuracy of the estimated Kinetic parameters through the following features:

1. the cumulative integral of all the surfaces that can exist between the estimated states trajectory and the simulated or known ground truth. We use the trapezoidal method with varying spacing which is particularly adapted to the shape of the idealized traces since the areas between the estimated state trajectory and the ground truth are rectangles, making the resulting approximations of such integration method more accurate (because the estimated and target idealized traces to integrate are both stepwise functions, the trapezoids are rectangles). Let *f* (*x*) be the estimated idealized trace and *g*(*x*) the ground truth states trajectory. We associate the residuals to all the areas between *f* and *g*, and discretized them into *N* non-uniform intervals. For each interval [*a, b*], we calculate the integral 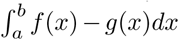 by partitioning the integration domain [*a, b*] into *m* sub-intervals *{x*_*k*_*}* such that *a* = *x*_0_ *< x*_1_ *< … < x*_*m−*1_ *< x*_*m*_ = *b* and Δ*x*_*k*_ = *x*_*k*_ − *x*_*k−*1_ the length of the *k*^*th*^ subinterval, then the cumulative integration of the residuals over the total number *r*_*max*_ of residual areas is: 

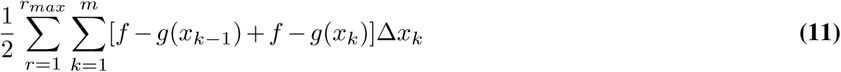

To better visualize such residual areas due to missed or fake transitions, and how well all the scenarios are captured into the proposed cumulative integral, we simulated a 5-states model and run cc-EM.

As explained before, the trapezoidal method used to integrate the different residuals areas due to the errors between the estimated idealized trace *f* (*x*) and the ground truth states trajectory *g*(*x*), splits the integration domains into several sub-intervals having rectangle shapes because *f* and *g* are stepwise functions.

While the integration domains are not uniform between the residual areas and depend on the estimated transitions, their corresponding sub-intervals all have a single data point resolution (bin width), as can be seen in figure 10 through the x-coordinates of the corners of the pink shaded residual area. By doing so, we manage to scan the whole estimated idealized trace data point by data point and to precisely localize and quantify all the transitions estimation errors. Not only such error integration method provides qualitative information about false positive (grey shaded areas) and false-negative transitions (pink shaded areas), by covering all the scenarios of the stepwise regression and states classification in both the signal magnitude (transition amplitude) and time dimensions (transition index), thanks to the rectangle shape of the integration domains, but also it informs directly on the possible biases of all the multi-state model parameters: states’ level, states’ population, transition index. The OKB metric considers also the total number of detected transitions (true and false) and normalizes the resulting residual areas by the sample size since the states and transitions detection error probability depends also on the time course trajectory of the stochastic process to model.

The OKB metric also relies on two other kinetic features for the penalization of the trace idealization objective:

2) The state-space size estimate with a greater weighting compared to the other features because a wrong molecular model is more serious than the kinetic parameters estimation errors of a model that correctly explains the phenomena under monitoring
3) The reaction pathway coordinates to check if the idealized trace correctly links the states chronologically, which is another important kinetic parameter especially for the mean first passage times (after how long is each state visited for the first time) of stochastic processes.

The color code proposed here to differentiate the estimation errors is to retrieve the qualitative information related to the false positive and false negative transitions. Finally, we have the following formula of the overall accuracy of the kinetic parameters estimate: 

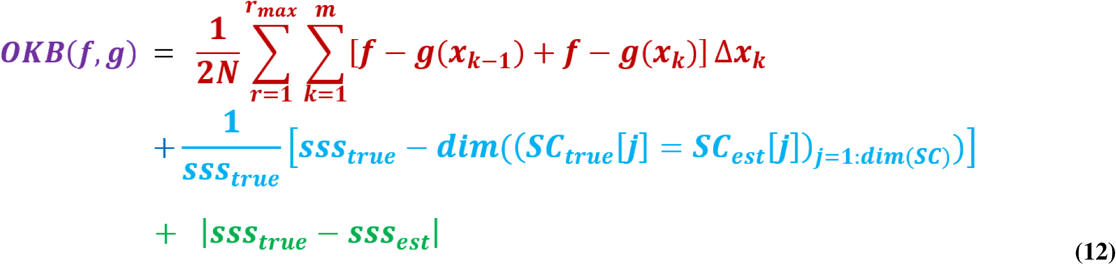

The first term in Eq.12 is the normalized cumulative integral of the residual areas induced by both missed and fake transitions in the estimated idealized trace. The second term corresponds to a penalty of the states’ coordinates in the estimated idealized trace compared to the ground truth, that is we extract from the idealized trace and the simulated states trajectory respectively two vectors *SC*_*est*_ and *SC*_*true*_ that contain the sequence of states in the chronological order in which they have been first visited within the sampling window. So that if each element of *SC*_*est*_ is equal to those of *SC*_*true*_, the resulting states coordinate penalty is null, while the upper bound of this penalty equals to 1 if we did not manage to retrieve the right chronological sequence of the visited states. The last term encodes for the state space size estimation error, because even if the idealized trace is very similar to the ground truth states trajectory, and the estimated states’ coordinates error close to zero, if the molecular model itself is wrong (i.e. if the estimated number of hidden states is different of the true expected number), whatever the underlying kinetic parameters estimates, we must penalize such misestimate given a higher weighting. All in all, the OKB score is a real positive valued metric and we can assess the performance modeling of the proposed algorithm according to the following scoring rules:

- if *OKB <* 0, we have an underfitting of the state space size.
- if *OKB* = 0 we have a perfect match between the idealized trace and the ground truth, given the right states coordinates along the reaction pathway, and the right state space size.
- if 0 *< OKB <* 1, represents an excellent level of overall accuracy which means that the estimated values of the kinetic parameters have a relative low bias compared to their true target values. Such range of OKB scores also guarantees both the right molecular model and states coordinates since the second and third terms of the OKB score are null.
- if *OKB ≥* 1, the proposed framework informs whether or not we have the right number of hidden states (in the case of simulated trace) and the right states’ coordinates. If so, an OKB score greater than one means that the idealized trace has a significant rate of inaccurate transitions, while the user can access to both the number and the type of misestimates (false positive and false negative states and transitions).

Finally, we provide in fig. 11 the OKB score landscape of the cc-EM algorithm across the two main limiting factors, the noise level and the number of states, to guide the user to reach the best outputs in terms of trace idealization and kinetic parameters estimates.

**Fig. 13.**
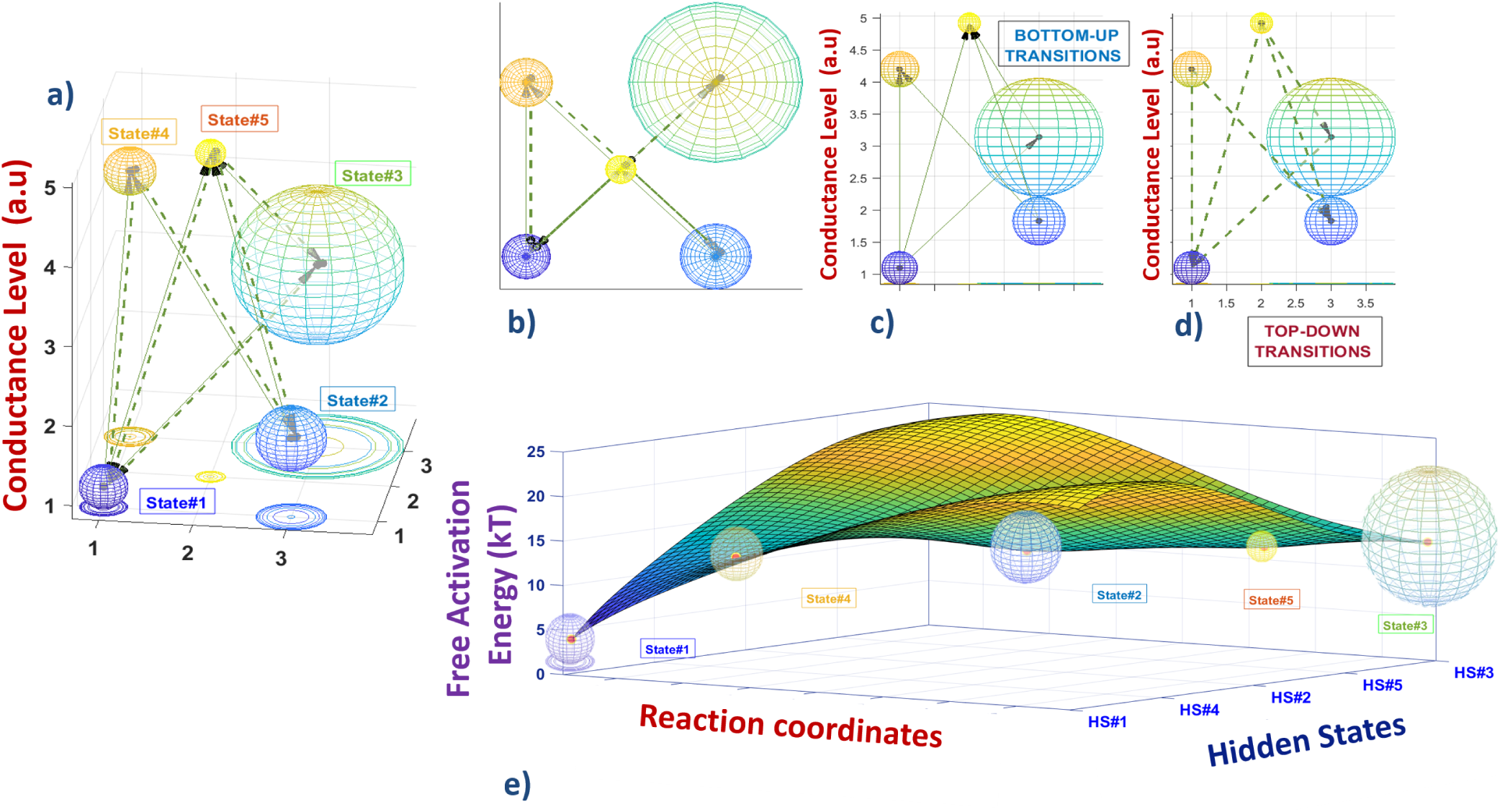
simoVIS data visualization of a simulated 5-states smFET signal: a) a 3D cartoon depicting the states and transitions network wrt the conductance levels of each sensor response to the conformational states. b) top view allowing to better highlight that actually some transitions between states are forbidden (for instance *state*#1 *↔ state*#2, *state*#2 *↔ state*#3, *state*#3 *↔ state*#4, *state*#4 *↔ state*#5). c) side view of the bottom-up transitions represented by solid line vectors between states having the lowest conductance level to the states with higher sensor response c) side view of the top-down transitions represented as dash vectors linking the states according to the descending conductance levels e) Free activation energy landscape of the reaction system with the hidden states as spheres of different occupancy rate, while the best path between the end-states can be clearly visualized state by state with the corresponding reaction coordinates *state*#1 *→ state*#4 *→ state*#2 *→ state*#5 *→ state*#3.

## Supplementary Note 3: Trace Idealization under Baseline Drift

We provide a built-in baseline drift removal tool with the trace idealization framework, users can refer to our previous work (41) detailing the signal and sensor parameters that have been tested to assess the robustness of the drift compensation algorithm. Basically, the MDL-AdaCHIP algorithm outperforms other baseline drift compensation methods, whatever if the parasitic baseline wander has the same signal characteristics or not as the useful signal, while being both non-stationary, and without any supervision nor signal prefiltering. Hereinafter we present a challenging case through a 15-states model with fast and numerous transitions and short-lived states, whose the states trajectory has been corrupted by a simulated sine baseline drift. In panel 12c), we clearly see how critical is the parasitic baseline drift as it introduces fakes states and transitions in the molecular dynamics time trajectory, and why ignoring drift can lead to erroneous outcomes of the trace idealization.

## Supplementary Note 4: simoVIS: a SIngle-MOlecule kinetics VISualization Tool

We proposed a companion tool to the cc-EM framework as an alternative single-molecule data visualization approach to the conventional representation of the states and transitions trajectory as a piecewise discrete signal, enabling to capture at a glance the kinetics of the molecular dynamics in an easier manner than the learned transition probabilities matrix. The simoVIS transition network does not correspond to the restricted states and transitions detected in the limited and finite acquired signal, but rather depicts the inferred states and rates beyond the sampling window through the learned transition matrix and its eigenvalue/eigenvector spectrum using 

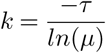

with *k* a rate, *τ* the lag time and *µ* an eigenvalue (67).

In the cartoon 13, each sphere represents a molecular state and the arrows correspond to the transitions between states, that is the kinetic changes in the reaction system. The radius of each sphere is proportional to the state occupancy rate inferred from the density population of the corresponding state, so that the more visited is a state, the biggest its corresponding sphere will be. Concerning the transitions, they are associated to vectors, while the data visualization tool offers two options: a first representation separating the incoming transitions to a state, from the state’s outgoing transitions, and a second alternative visualization separating the orientation and the sense of the transitions with respect to the intensity level of the involved states.The former representation is a visual representation of the transition probabilities matrix, while the latter representation takes into account the transition amplitudes to figure out the energy landscape of the states. In both cases, the width of the arrows is proportional to the transition rates, so that the more frequent are the exchanges between states, the thicker the transition vectors are. simoVIZ also provides the reaction pathway through the states’ coordinates and the corresponding mean first passage times (in case of Markovian processes), while the Gibbs energy landscape positions the local energy minima of each sub-states and end-states.

**Table 2.**
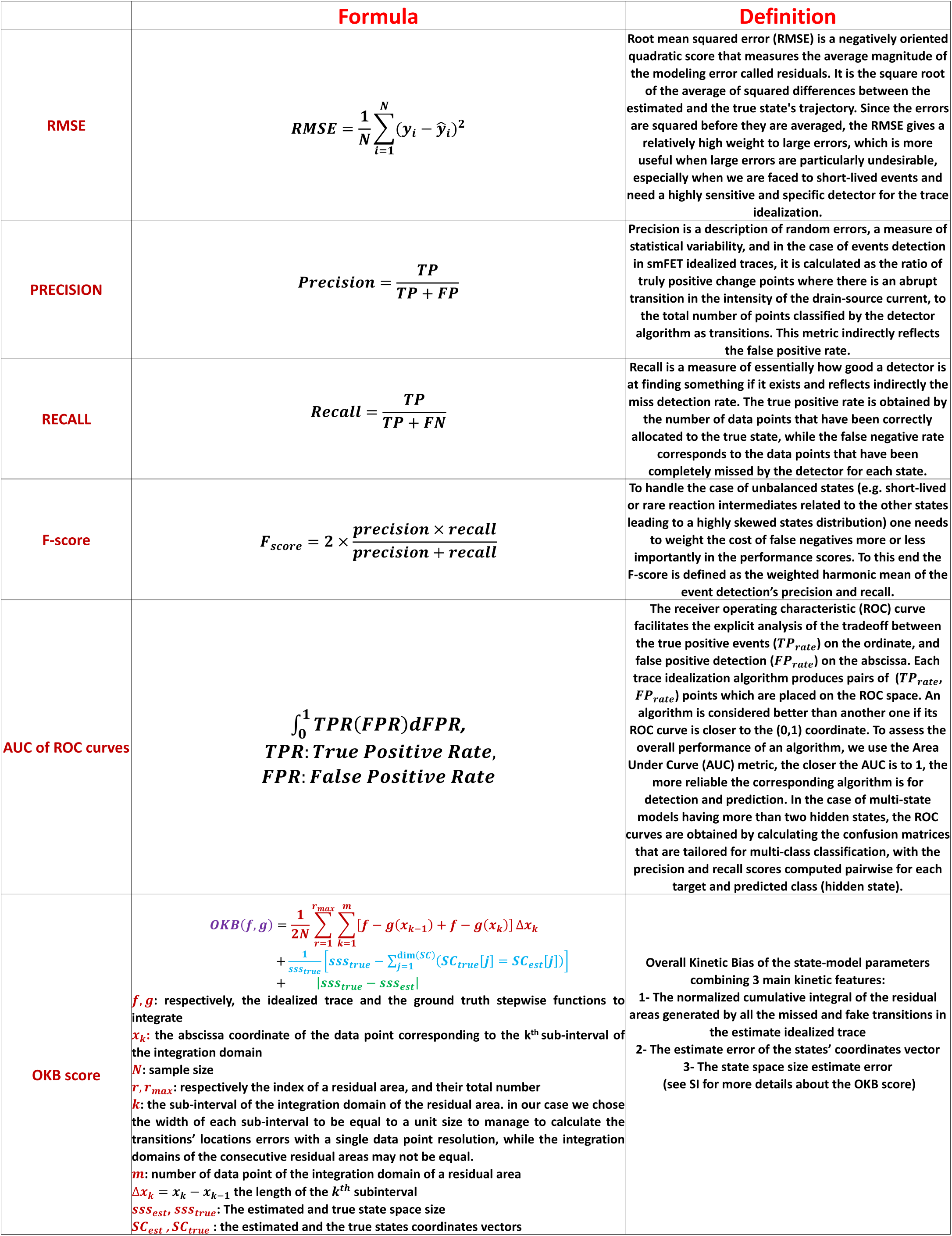
Performance metrics for the validation tests of cc-EM

